# Altered auditory maturation in Fragile X syndrome and its involvement in audiogenic seizure susceptibility

**DOI:** 10.1101/2025.04.22.650065

**Authors:** Dorit Möhrle, Demi Ma, Wenyue Xue, Jun Yan, Ning Cheng

## Abstract

**Background:** Auditory hypersensitivity is a prominent symptom in Fragile X syndrome (FXS), the most prevalent monogenic cause of autism and intellectual disability. FXS arises through the loss of the protein encoded by the *FMR1* (Fragile X Messenger Ribonucleoprotein 1) gene, FMRP, required for normal neural circuit excitability. In the brainstem, FMRP is necessary for normal development of acoustic reactivity, and its loss has been implicated in audiogenic seizures (AGS) in *Fmr1* knockout (KO) mice, modelling auditory hypersensitivity and seizures in FXS patients.

**Purpose:** The present study investigated the correlation between auditory brainstem function and behavioral expression of AGS at the early (postnatal day P20, infancy) and late (P32, juvenile) stage of auditory development in *Fmr1* KO mice compared with wildtype (WT) mice, and in both females and males.

**Methods:** We tested responsiveness to pure tones of select auditory pathway elements through auditory brainstem responses; and neural synchronization to amplitude envelopes of modulated acoustic stimuli through auditory steady-state responses. AGS behavior was categorized for severity during 5-minute exposure to loud sound. Expression of the immediate early gene cFos was quantified as a marker for neuronal activity in the inferior colliculus.

**Results:** During infancy, more severe AGS expression in *Fmr1* KO mice compared with WT mice was accompanied by increased responsiveness to acoustic stimuli at the level of the superior olivary complex and inferior colliculus, and stronger neural synchronicity in subcortical auditory neurons. *Fmr1* KO mice also had higher cFos positive cell counts in the inferior colliculus after exposure to loud sound. With age, both AGS susceptibility and exaggerated acoustic stimulus-evoked activity in the *Fmr1* KO mice subsided. Intriguingly, *Fmr1* KO mice displayed altered developmental profile in both the threshold and amplitude of auditory brainstem response.

**Conclusion:** Our findings support evidence that AGS activity relies upon hyperexcitability in the auditory system, including in the lower brainstem, possibly due to disturbed auditory maturation. Hyper-synchronization to modulated sounds in subcortical auditory neurons seemed to predict AGS severity. A better understanding of FXS-related circuit and behavioral symptoms of auditory processing across development provides the potential to identify therapeutic strategies to achieve auditory function recovery in FXS.

## 1. Introduction

The brain’s ability to process sounds undergoes a tremendous amount of development in early life so that we may hear and communicate effectively. During early postnatal development, the auditory system is particularly sensitive to sensory input during specific time windows of enhanced plasticity, with these windows varying depending on the complexity of the sounds being processed (Anbuhl et al., 2022; Buran et al., 2014; Keuroghlian and Knudsen, 2007; Kral, 2013a). For instance, critical periods for simple sound features, such as pure frequency tones, occur earlier, whereas more complex sound features undergo their own distinct periods of plasticity later in development (Barkat et al., 2011; Bhumika et al., 2019; de Villers-Sidani et al., 2007; Nakamura et al., 2020). This sensitive period-driven development allows experience to shape the auditory system and its functional capabilities, but disruptions during these periods can have negative, long-lasting effects on auditory processing (Keuroghlian and Knudsen, 2007; Kral, 2013b). This auditory maturation is commonly altered in neurodevelopmental disorders, such as autism and Fragile X syndrome (FXS, Bhumika et al., 2019; Contractor et al., 2015; LeBlanc and Fagiolini, 2011; Takesian and Hensch, 2013), which can ultimately lead to auditory processing difficulties (Sinclair et al., 2017).

FXS, the leading inherited cause of autism and intellectual disability, is caused by a trinucleotide repeat expansion in the *Fragile X Messenger Ribonucleoprotein 1* (*Fmr1*) gene, resulting in the loss of Fragile X Messenger Ribonucleoprotein (FMRP) expression (Fu et al., 1991; Hagerman et al., 2008; Warren and Nelson, 1994). A hallmark of both autism and FXS is sensory hypersensitivity and hyperreactivity. One of the most prevalent, persistent, and disabling sensory challenges of FXS and autism is auditory hypersensitivity, where a majority of affected individuals perceive everyday sounds as unbearably loud and show reduced tolerance to sound (Gomes et al., 2008; Rais et al., 2018; Sinclair et al., 2017; Williams et al., 2021a; Williams et al., 2021b). These challenges are associated with elevated cortical responses and decreased ability to habituate the neural response to repeated sounds (Castrén et al., 2003; Ethridge et al., 2016; Knoth et al., 2014; Wang et al., 2017). The *Fmr1* knockout (KO) mouse, a well-established model for FXS and syndromic autism, recapitulates the human auditory hypersensitivity phenotype in both behavioural and electrical responses (Rotschafer and Razak, 2014): *Fmr1* KO mice show hyperreactivity to moderately intense prepulse tones, as evidenced by an increased suppression of the startle response in the prepulse inhibition of the acoustic startle response paradigm. Additionally, they are susceptible to audiogenic seizures (AGS) induced by high-intensity acoustic stimulation (Chen and Toth, 2001; Musumeci et al., 2000; Rotschafer and Razak, 2013). AGS in *Fmr1* KO mice involve the ascending auditory system, including the central auditory pathway, and are known to depend on the brainstem, i.e., the inferior colliculus (IC) in the midbrain (Fig. 1C) (Chen and Toth, 2001; Gonzalez et al., 2019; Lovelace et al., 2018; Nguyen et al., 2020). Neuronal hyperresponsiveness in the auditory cortex (AC) and IC, thought to be associated with behavioural hypersensitivity to sound stimuli in *Fmr1* KO mice, appears to emerge between the second and third postnatal week during early auditory development (Nguyen et al., 2020; Wen et al., 2018). Intriguingly, AGS in *Fmr1* KO mice exhibit a peak incidence between postnatal days P20 and P30, with particularly heightened susceptibility observed at the beginning of this timeframe (Dölen et al., 2007; Michalon et al., 2012; Musumeci et al., 2000; Nguyen et al., 2020; Pacey et al., 2009; Yan et al., 2005). It has been suggested that lack of FMRP, which is typically highly expressed in the auditory brainstem, leads to dysregulation of critical developmental windows in brainstem circuits and possibly transient disruptions in auditory brainstem signaling, resulting in this age-dependent hypersensitivity (Meredith et al., 2012; Wang et al., 2014; Yun et al., 2006; Zorio et al., 2017). Despite auditory hypersensitivity being a common and impactful symptom of FXS and other neurodevelopmental disorders, we still lack a clear understanding of the neural mechanisms and temporal maturation dynamics in the auditory brainstem that might contribute to this condition, and this has ultimately hindered efforts to support the affected individuals and families effectively.

**Fig. 1.**
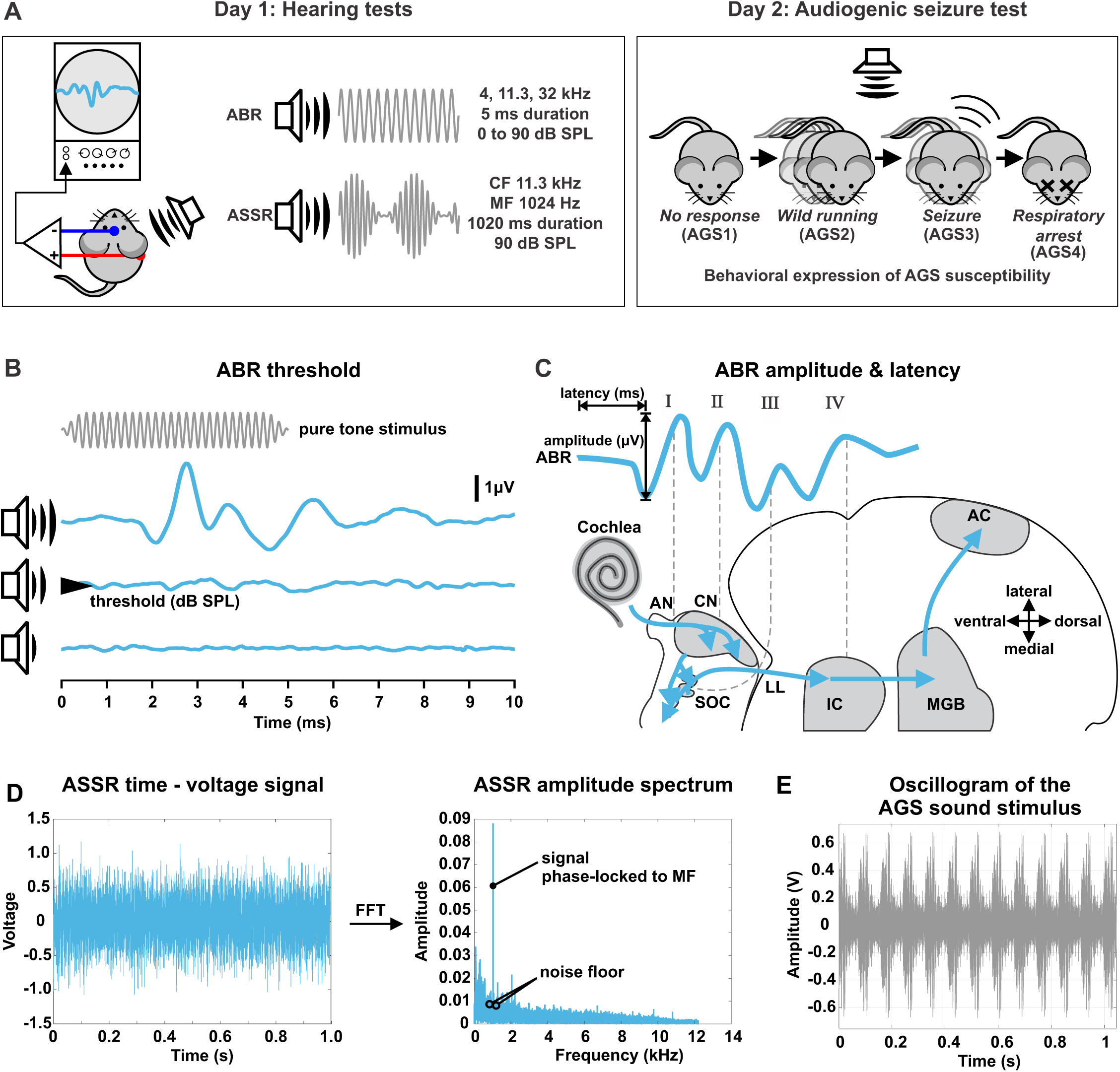
Experimental approach. **(A)** Timeline including hearing tests (ABR and ASSR measurements) and behavioral expression of AGS susceptibility. **(B)** Scheme of ABR threshold determination. **(C)** Schematic drawing of the auditory pathway and correlated stimulus-evoked deflections of ABR waves I, II, III, and IV. **(D)** Scheme of ASSR signal-to-noise extraction. **(E)** AGS test stimulus was an amplitude- and frequency modulated sound (mean frequency 4.8 kHz, amplitude modulation frequency ∼12 Hz). Abbreviations: ABR, Auditory Brainstem Response; AC, auditory cortex; AGS, AGS; AGS1, no response; AGS2, wild running; AGS3, tonic-clonic seizure; AGS4, respiratory arrest; AN, auditory nerve; ASSR, Auditory Steady State Response; CF, Carrier frequency; CN, cochlear nucleus; FFT, Fast Fourier Transformation; IC, inferior colliculus; LL, lateral lemniscus; MF, Modulation frequency; MGB, medial geniculate body; SOC, superior olivary complex.

In the present study, we investigated the correlation between auditory brainstem function and the behavioral expression of AGS, used as a proxy for auditory hypersensitivity, in *Fmr1* KO mice at two stages of auditory development: early (P18-20, infancy) and late (P31-34, juvenile). We compared behavioral and electrophysiological responses of *Fmr1* KO mice to age-matched wildtype (WT) controls, allowing us to assess the effect of altered auditory maturation on brainstem processing and hypersensitivity. We tested both female and male mice of both the genotypes. Auditory function was assessed through auditory brainstem response (ABR) wave amplitudes and latencies, corresponding to neural responsiveness and transmission speed of distinct parts of the ascending auditory pathway: the auditory nerve (AN; wave I), the cochlear nucleus (CN; wave II), the superior olivary complex (SOC, wave III), and the lateral lemniscus (LL) and IC (wave IV, Fig. 1A, C) (Melcher and Kiang, 1996). Furthermore, we measured auditory steady-state responses (ASSRs) evoked by neural synchronization to the amplitude envelopes of spectrotemporally complex stimuli, in order to assess the synchronous, phase-locked discharge of subcortical auditory neurons to the modulation frequency of acoustic stimuli (Fig. 1A, D) (Kuwada et al., 2002; Parthasarathy and Bartlett, 2012; Picton et al., 2003). We found that hyperresponsiveness in the SOC and LL and IC of *Fmr1* KO mice at P20 was associated with a higher propensity for AGS. This aligned with the cessation of both seizures and hyperresponsiveness by P32. At P20, increased ASSRs to amplitude-modulated stimuli in *Fmr1* KO mice also correlated with heightened AGS severity. These results support evidence that AGS susceptibility relies upon hyperexcitability in the auditory system, including the lower brainstem, possibly due to disrupted maturation during early postnatal development. This temporal pattern highlights that early disruptions in auditory processing during this sensitive period may predispose individuals to auditory hypersensitivity and may indicate a time window when the brain is more susceptible to therapeutic treatments.

## 2. Materials and Methods

### 2.1. Mice

Breeder FVB wildtype (WT, FVB.129P2-Pde6b^+^ Tyr^c-ch^/AntJ, strain #:004828) and *Fmr1* KO mice (FVB.129P2-*Pde6b*^+^ *Tyr^c-ch^ Fmr1^tm1Cgr^*/J, strain #:004624) were obtained from the Jackson Laboratory (ME) and the lines were maintained at the mouse facility of the Clara Christie Centre for Mouse Genomics (CCCMG), University of Calgary. Mice were group-housed with up to five mice per cage in a humidity- and temperature-controlled room with a 12-h light/dark cycle and were fed standard mouse chow *ad libitum*. Hearing measurements were performed at either postnatal day P18 to 19 for infants or P31 to 33 for juveniles. Behavioral AGS expression was assessed one day after the hearing measurements (Fig. 1A). In the following, the age of infants is termed P20 and the age of juveniles P32. Four experimental groups were tested: WT_P20 (females F=9, males M=7), *Fmr1* KO_P20 (F=7, M=8), WT_P32 (F=6, M=4), and *Fmr1* KO_P32 (F=6, M=6). All procedures in this study were performed in accordance with the recommendations in the Canadian Council for Animal Care. The protocol of this study was approved by the Health Sciences Animal Care Committee of the University of Calgary.

### 2.2. Hearing Measurements

ABRs and ASSRs were recorded in anaesthetized mice (Fig. 1A, infants: 50 mg/kg ketamine hydrochloride, 2.5 mg/kg xylazine hydrochloride, juveniles: 100 mg/kg ketamine hydrochloride, 5 mg/kg xylazine hydrochloride intraperitoneal injections) in a soundproof chamber similar to (Marchetta et al., 2020; Möhrle et al., 2016; Möhrle et al., 2017; Wolter et al., 2018) with subdermal silver wire electrodes at the ipsilateral (right) mastoid and the vertex of the animals. Anesthesia was supplemented if needed. All tests were conducted using a Tucker Davis Technology System 3 with Real-time processors RP2 and controlled with Jan Schnupp’s BrainWare software (V9.21 for TDT). None of the animals showed evidence of middle ear pathology or obstructive cerumen. Body temperature was maintained at 37°C during testing through a heating pad in the experimental chamber. All recordings were blinded for the subjects’ experimental conditions before analysis using custom-made MATLAB scripts (R2022a, The MathWorks).

#### Auditory brainstem response (ABR)

After amplification (50,000-fold) and filtering (300-3000 Hz) (Grass Telefactor P55 AC preamplifier, line filter on), the ABR signals were averaged for 512 repetitions at each sound pressure presented (0–90 dB SPL in steps of 5 dB). ABR thresholds were determined with pure tone stimuli (5 ms, including cosine-gated 0.5 ms rise/fall time, 4, 11.31, and 32 kHz) as the lowest sound pressure that produces visually distinct evoked potentials (Fig. 1B) by an expert observer.

ABR waveforms were analyzed for consecutive amplitude deflections (waves, Fig. 1C), with each wave consisting of a starting negative peak and the following positive peak. Peak amplitudes and latencies of ABR waves I, II, III, and IV were extracted using custom-made MATLAB scripts. Wave latencies were defined by the onset timing (negative peak) of each corresponding wave and ABR wave amplitudes as the peak-to-peak difference. From the extracted peaks, ABR peak-to-peak (wave) amplitude and latency growth functions were calculated for individual animals for increasing stimulus levels. All ABR wave amplitude and latency growth functions were normalized with reference to the ABR thresholds. To analyze the slopes of the ABR wave amplitude growth functions in the dB range with rapid growth, we calculated the 99% confidence interval for each individual recording and extracted the response range that continuously exceeded this interval. We used linear regression to compare the slopes between experimental groups.

#### Auditory steady state response (ASSR)

ASSRs were elicited with amplitude-modulated sinusoidal stimuli (1020 ms, including cosine-gated 5 ms rise/fall time) using a 11.31 kHz carrier and modulation frequency of 512 Hz at 90 dB SPL and 100% modulation depth. The signals were filtered (100-10000 Hz) and averaged for 192 repetitions to compute the ASSR time–domain waveform (Fig. 1D). Fast Fourier transforms (FFTs) were performed on the waveforms starting 20 ms after stimulus onset and ending 25 ms before stimulus offset to exclude transient ABRs and offset responses using custom-written programs in MATLAB. To calculate the ASSR signal-to-noise-ratio (SNR), we extracted the maximum amplitude at 1024 Hz (i.e., the first harmonic of the modulation frequency) as well as the amplitude of the noise floor and divided these two values (Fig. 1D). In more detail, to determine the amplitude of the FFT peak signal, we examined the frequency bin at 1024 Hz, along with the two adjacent bins above and below it (each 1 Hz wide). The maximum value within these three bins was defined as the signal amplitude. To estimate the noise floor, we excluded the six bins directly adjacent to the signal (three bins above and three bins below), and then averaged the amplitudes from the two three-bin intervals adjacent to these excluded bins. The ASSR at the first harmonic of the amplitude modulation frequency is reported as the FFT SNR.

### 2.3. AGS testing

To evaluate AGS, mice were placed in a plastic chamber (42 × 42 × 42 cm) which was closed with a fabric covered lid containing a ∼90dB siren. The siren sound (Fig. 1E) was presented to mice immediately for a duration of 5 min. Mice were categorized for behavioral phenotype as described based on experimenter observation: AGS1 = no response, AGS2 = wild running, AGS3 = tonic-clonic seizures, AGS4 = respiratory arrest or death as previously described (Dölen et al., 2007; Ronesi et al., 2012). Mice were given at least 1 h to acclimate to the test room before the behavioral experiment.

### 2.4. Tissue preparation and immunohistochemistry

We quantified the expression of the immediate early gene cFos as a marker for neuronal activity in the auditory system (Ehret and Fischer, 1991) in a separate cohort of WT_P20 (F=3, M=3), *Fmr1* KO_P20 (F=4, M=3), WT_P32 (F=3, M=3), and *Fmr1* KO_P32 (F=3, M=3) mice. The animals were exposed to the AGS test siren for 5 min, and then were deeply anesthetized. The mice were perfused with ice-cold 0.1 M PBS, pH 7.4 (1X PBS) followed by 4% paraformaldehyde (PFA) in ddH2O (∼15 min after sound exposure). Brains were extracted, post-fixed for 48 h in 4% PFA at 4°C, and transferred to 30% sucrose in 1X PBS.

Coronal sections (40 µm) containing the IC were cut on a cryostat (Leica CM1850 UV) and kept in 0.02% sodium azide (MKBL4701V, Sigma-Aldrich) in 1X PBS. After washing three times in Tris-buffered saline with 0.1% Tween®20 detergent (1X TBST; 1, 5, 10 minutes), the sections were incubated in blocking buffer containing 50 mM glycine in 1X TBS containing 0.1% gelatin, 5% NHS and 0.5% Triton-X 100 for 90 minutes at room temperature, and then with primary antibody (c-Fos (9F6) rabbit mAb #2250, Cell Signalling Technology, 1:1000) in antibody solution (0.1% hydrogen peroxide, 0.2% sodium azide, 10 mM glycine in 3% NGS and 0.2% Triton-X100 in 1X TBS) for three days at 4°C. After washing three times with 1X TBST (1, 5, 10 minutes), the sections were incubated with secondary antibody (Cy™3-conjugated AffiniPure™ donkey anti-rabbit IgG (H+L), Jackson ImmunoResearch Laboratories, INC., 1:2000) and 4′,6-diamidino-2-phenylindole (DAPI, 1:50) in antibody solution (0.1% hydrogen peroxide, 10 mM glycine in 3% NGS and 0.2% Triton-X100 in 1X TBS) at room temperature for 2 h. After another three wash cycles in 1X TBST (1, 5, 10 minutes), sections were mounted with Fluoromount-G® medium (Cat. No.: 0100-01, SouthernBiotech) on microscope slides (MSSF-50, FroggaBio) and cover slipped (MCG-100, FroggaBio). Fluorescent images from the left and right central IC were collected for each animal using an epifluorescence microscope (Olympus BX61) connected to a camera (FLIR Blackfly® S BFS-U3-16S2M 20523795) and an image capturing software (Spinnaker^®^SDK SpinView 1.10.0.31). We used the open-source ImageJ/Fiji tool ‘Quanty-cFos’ (Beretta et al., 2023) to detect cFos positive cells in the microscopic images and derive of cell counts (StarDist 2D, Batch Analysis, intensity cut-off: automatic, area cut-off: 160, with manual optimization). In more detail, to overcome variability in differentiating cFos positive and negative cells as a result of image brightness, we opted for automated intensity threshold detection in ‘Quanty-cFos’. As demonstrated by Beretta et al. (2023) this approach is superior to manually counting cells in that it results in more consistent cFos positive cell counts. The area threshold was initially determined through trial and error to identify the most accurate value for detecting cFos-positive cells and by comparing results from three manual counters. Using an area threshold value of 160, almost all cFos positive cells were consistently identified across a number of sample images, with the occasional 5 or fewer cells that had to be added if false negative or removed if false positive. After using the tool to detect cFos postive cells, all images were reviewed to ensure correct cell count and the counts were edited if necessary. The cell counts from the left and right IC of each animal were averaged before conducting comparisons between experimental groups.

### 2.5. Statistical analysis

Statistical tests were performed in GraphPad Prism 9.5.1 (GraphPad Software, San Diego, California) and RStudio 2022.2.0.0 (PBC, Boston, Massachusetts), and figures were generated in GraphPad Prism and in CorelDRAW X7 (Alludo, Ottawa, Ontario, Canada). Data following normal distribution (D’Agostino & Pearson test or Shapiro-Wilk test for small sample sizes) were analyzed using unpaired *t* test, or ordinary one-way ANOVA and *post hoc* multiple comparison tests with correction for type 1 error after Dunnett’s method. To accommodate for non-normal distribution and missing values we used ARTool (Aligned Rank Transform, ART) to align-and-rank data for nonparametric two-way, three-way, and mixed effects (repeated measures) ANOVA for main effects and interactions (Wobbrock et al., 2011), and ART-C for *post hoc* pairwise comparisons (contrast tests, Elkin et al., 2021). All tests included the effect of sex. *Post hoc* tests (paired for dB *re* threshold) for statistically significant interactions were not significant unless stated otherwise. As a measure for effect sizes, we calculated partial eta squared *ηp²* for ART ANOVAs, estimates of the difference and standard error for ART-C tests, and the mean difference and 95% confidence interval for *t* tests and Dunnett’s tests. Correlations were tested using Pearson’s correlation coefficient for continuous data or Spearman’s correlation coefficient for categorized data (r value <0.1, negligible; 0.1-0.4, weak; 0.4-0.7, moderate; 0.7-0.9, strong; 0.9-1.0, very strong correlation, Rowntree, 2004) and *p* value to quantify the likelihood of a correlation. Simple logistic regression was used for predicting binary outcomes of the AGS test. Statistical significance level was *α* = 0.05, and resulting *p* values are reported in the figure legends and tables using: \**p* < 0.05; \*\**p* < 0.01; *p*\*\*\* < 0.001. Data are presented as group mean and standard error of the mean (SEM) or as group median and interquartile range (IQR) as stated in the figure legends. Individual data points represent individual animals.

## 3. Results

### 3.1. AGS susceptibility in *Fmr1* KO mice decreases with age

To determine the effect of auditory maturation on sensory hypersensitivity in FXS, we categorized AGS in both infant (P20) and juvenile (P32) WT and *Fmr1* KO mice. AGS susceptibility was examined by observing the behavioral responses during 5-minute exposure to loud sound (Fig. 1A, E). All WT_P20 mice showed no response (100%, Fig. 2A) and we observed typical exploratory and self-grooming behaviors. In comparison, less *Fmr1* KO_P20 mice exhibited no response (27%). Consistent with previous findings (Gonzalez et al., 2019; Sawicka et al., 2016; Wang et al., 2012), *Fmr1* KO_P20 mice showed robust AGS phenotypes, with 20% wild running, 20% tonic-clonic seizure, and 33% respiratory arrest (Fig. 2A). In contrast to infants, both juvenile WT_P32 and *Fmr1* KO_P32 mice had no response (100%, Fig. 2B). These findings indicate that *Fmr1* mutation leads to robust AGS expression during a time that coincides with early auditory development. By the time of late auditory development, AGS susceptibility had subsided.

**Fig. 2.**
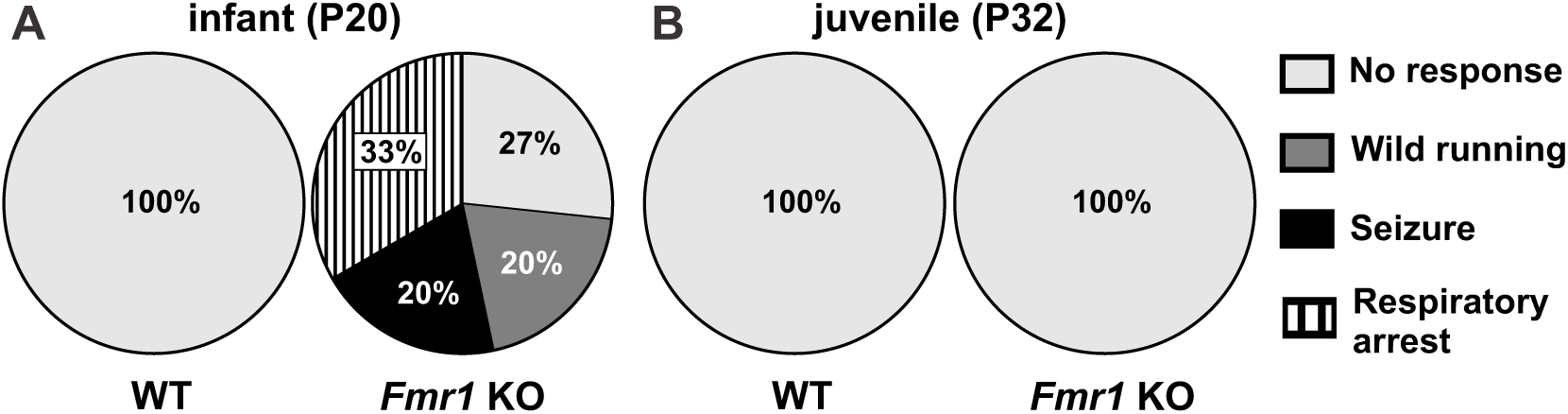
AGS susceptibility in *Fmr1* KO mice decreased with age. **(A)** During infancy, *Fmr1* KO mice, but not WT mice, displayed a range of behavioral severity in response to loud sound during AGS testing. **(B)** Both juvenile WT and *Fmr1* KO mice show no behavioral response to loud sound. Legend applies to **(A)** and **(B)**: no response (consisting of pause or continuous exploration), wild running, seizure, or respiratory arrest (death). Infants (P20), WT (*n*=12), *Fmr1* KO (*n*=16); juveniles (P32), WT (*n*=10), *Fmr1* KO (*n*=12). Data expressed as percent of animals.

### 3.2. *Fmr1* deletion delays maturation of auditory detection thresholds

Previous studies suggest a relationship between hearing deficits and AGS severity (Faingold et al., 1990; Ralls, 1967). We estimated basic hearing function of infant and juvenile WT and *Fmr1* KO mice using ABR measurements (Fig. 1A). We stimulated the auditory system along low-to-high frequency hearing range with pure tone stimuli that allow for the frequency specific allocation of auditory responses to a tonotopic place in the cochlea of the auditory periphery (Müller, 1991).

First, we compared ABR thresholds (Fig. 1B) to determine if *Fmr1* mutation led to altered maturation of hearing sensitivity. As expected (Song et al., 2006; Zhou et al., 2006), ABR thresholds varied depending on stimulus frequency (Fig. 3, Supplementary Table 1). In general, thresholds at 11.3 kHz were the lowest (mean±SEM, 22.5±0.82 dB) and those at 32 kHz (33.4±1.19 dB) were lower than at 4 kHz (52.5±0.90 dB; Fig. 3, Supplementary Table 2). This means that the hearing was the most sensitive around the 11.3 kHz region, as is typical for mice (Ehret, 1974; Heffner and Heffner, 2007; Ralls, 1967). The ABR thresholds at all three frequencies decreased with age (Fig. 3, Table 1). A significant influence of genotype was observed only at 32 kHz (Table 1). In particular, ABR thresholds in *Fmr1* KO_P20 mice (41.1±1.9 dB) were higher than in WT_P20 mice (34.4±1.76 dB), whereas those in *Fmr1* KO_P32 mice (29.6±1.3 dB) were similar to WT_P32 mice (25.5±2.29 dB; Fig. 3, Table 2).

**Fig. 3.**
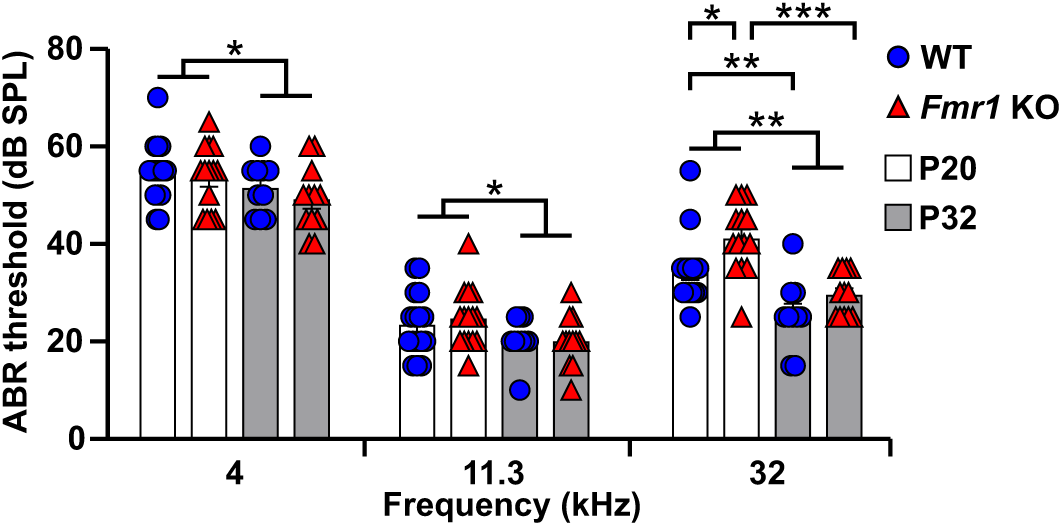
*Fmr1* knock-out delayed maturation of hearing sensitivity in the high frequency range. Hearing thresholds from infant (P20, white bars) and juvenile (P32, gray bars) WT (blue circles) and *Fmr1* KO mice (red triangles) assessed from electrical ABR potentials in response to low, medium, and high frequency pure tone auditory stimuli (4, 11.3, and 32 kHz). ABR thresholds decreased with age independent of genotype. Additionally, at 32 kHz, ABR thresholds were higher in *Fmr1* KO_P20 compared with WT_P20 mice. 4kHz and 32kHz, WT_P20 (*n*=16), *Fmr1* KO_P20 (*n*=14), WT_P32 (*n*=10), and *Fmr1 KO_P32* (*n*=12); 11.3kHz, WT_P20 (*n*=16), *Fmr1* KO_P20 (*n*=15), WT_P32 (*n*=10), and *Fmr1 KO_P32* (*n*=12). Data expressed as mean (bars) ± SEM (error bars) and individual animals (symbols). *p* values, *** *p* < 0.001, \*\**p* < 0.01 and * *p* < 0.05.

**Table 1.** Statistical comparisons of ABR thresholds from mice of the following four groups: WT_P20 (*n*=16), *Fmr1* KO_P20 (*n*=14-15), WT_P32 (*n*=10), and *Fmr1 KO_P32* (*n*=12). The thresholds for one *Fmr1* KO_P20 animal at 4 and 32 kHz were excluded from analysis because the acoustic stimuli were presented in 10 dB instead of 5 dB steps. This table is associated with data in Fig. 3.

| Parameter | Source of Variation | P value | P value summary | F (DFn, DFd) | $\eta p^2$ |
| --- | --- | --- | --- | --- | --- |
| ABR threshold at 4kHz | Genotype | 0.24 | ns | F(1, 48) = 1.35 | 0.027 |
|  | Age | 0.02 | * | F(1, 48) = 5.43 | 0.102 |
|  | Genotype:Age | 0.54 | ns | F(1, 48) = 0.37 | 0.008 |
| ABR threshold at 11kHz | Genotype | 0.90 | ns | F(1, 49) = 0.01 | <0.001 |
|  | Age | 0.04 | * | F(1, 49) = 4.10 | 0.077 |
|  | Genotype:Age | 0.46 | ns | F(1, 49) = 0.53 | 0.011 |
| ABR threshold at 32kHz | Genotype | 0.007 | ** | F(1, 48) = 7.69 | 0.138 |
|  | Age | <0.001 | *** | F(1, 48) = 28.83 | 0.375 |
|  | Genotype:Age | 0.21 | ns | F(1, 48) = 1.59 | 0.032 |

Two-way ART ANOVA, $p$ values, \* $p < 0.05$ , \*\* $p < 0.01$ , \*\*\* $p < 0.001$ , ns not significant.

**Table 2.** *Post hoc* pairwise comparisons for ABR threshold at 32kHz from mice of the following four groups: WT_P20 (*n*=16), *Fmr1* KO_P20 (*n*=14), WT_P32 (*n*=10), and *Fmr1 KO_P32* (*n*=12). This table is associated with data in Fig. 3.

| Parameter | Contrast | Estimate | SE | DF | t | Adjusted P Value | P value summary |
| --- | --- | --- | --- | --- | --- | --- | --- |
| ABR threshold at 32kHz | WT_P32 - WT_P20 | -15.66 | 4.51 | 48 | -3.47 | 0.005 | ** |
|  | WT_P32 - *Fmr1* KO_P32 | -6.63 | 4.79 | 48 | -1.38 | 0.51 | ns |
|  | WT_P32 - *Fmr1* KO_P20 | -27.50 | 4.63 | 48 | -5.94 | <0.001 | *** |
|  | WT_P20 - *Fmr1* KO_P32 | 9.03 | 4.27 | 48 | 2.11 | 0.16 | ns |
|  | WT_P20 - *Fmr1* KO_P20 | -11.84 | 4.10 | 48 | -2.89 | 0.02 | * |
|  | *Fmr1* KO_P32 - *Fmr1* KO_P20 | -20.88 | 4.40 | 48 | -4.74 | <0.001 | *** |

ART-C Tukey's multiple comparisons test, $p$ values, \* $p < 0.05$ , \*\* $p < 0.01$ , \*\*\* $p < 0.001$ , ns not significant.

Hearing sensitivity for high frequencies increases later during early postnatal development than for low or middle frequencies (Ehret, 1976). Therefore, the higher ABR thresholds at 32 kHz in *Fmr1* KO_P20 mice indicate that *Fmr1* mutation delayed the maturation of auditory detection thresholds in a frequency-specific manner. However, there was no correlation between ABR thresholds and AGS severity in *Fmr1* KO_P20 mice at any of the three stimulus frequencies (Pearson, 4 kHz: r 0.1485, *p* = 0.51; 11.3 kHz: r 0.2224, *p* = 0.44; 32 kHz: r 0.1693, *p* = 0.54). This suggests that the delayed maturation in hearing sensitivity might not directly participate in AGS susceptibility.

### 3.3. *Fmr1* deletion does not affect maturation of neuronal transmission speed along the auditory pathway

We analyzed above-threshold ABR wave level-latency and -amplitude functions corresponding to sound-evoked neuronal activity in the AN (wave I), in the CN (wave II), in the SOC (wave III), and in the LL and IC (wave IV; Fig. 1C, for exemplary ABR waveforms see Supplementary Fig. 1) (Melcher and Kiang, 1996). We focused the interpretation of differences in above-threshold hearing function of *Fmr1* KO_P20, WT_P20, *Fmr1* KO_P32, and WT_P32 mice in the middle frequency range (i.e., 11.3 kHz) because measurement variances were generally smaller for both genotypes. Since elevated ABR thresholds contribute to differences in latencies and amplitudes by reducing the effective stimulus levels, we expressed this analysis as dB *re* ABR threshold (i.e., ABR threshold is 0 dB *re* threshold). ABR wave latencies are known to mature rapidly over the course of the second and third postnatal weeks and to achieve adultlike characteristics thereafter (Song et al., 2006). As expected, ABR wave latencies were inversely related to sound level, and latencies of later waves were sequentially delayed relative to earlier waves (Fig. 4). In WT mice, ABR wave latency growth functions were significantly elevated in WT_P20 compared with WT_P32 mice across the auditory periphery to midbrain (wave I to IV, Fig. 4A, Table 3). In line with previous findings (Song et al., 2006), the range over which ABR wave latency values decreased during maturation was larger for the later than for earlier occurring waves. Specifically, on average, wave I latencies decreased by 0.22 ms, wave II by 0.28 ms, wave III by 0.44 ms, and wave IV latencies by 1.08 ms (across 0 to 65 dB *re* threshold). Similar developmental changes were observed in *Fmr1* KO mice (Fig 4B, Table 4): *Fmr1* KO_P20 mice showed significantly longer ABR wave I to IV latencies compared with *Fmr1* KO_P32 mice. Particularly, during maturation, wave I latencies decreased by 0.25 ms, wave II by 0.34 ms, wave III by 0.45 ms, and wave IV latencies by 1.08 ms (across 0 to 70 dB *re* threshold). Interestingly, for ABR wave IV in *Fmr1* KO mice, this developmental decrease in latency was particularly pronounced at near-threshold to lower sound levels (females: 10 to 40 dB *re* threshold, males: 0 to 35 dB *re* threshold; Table 5).

**Fig. 4.**
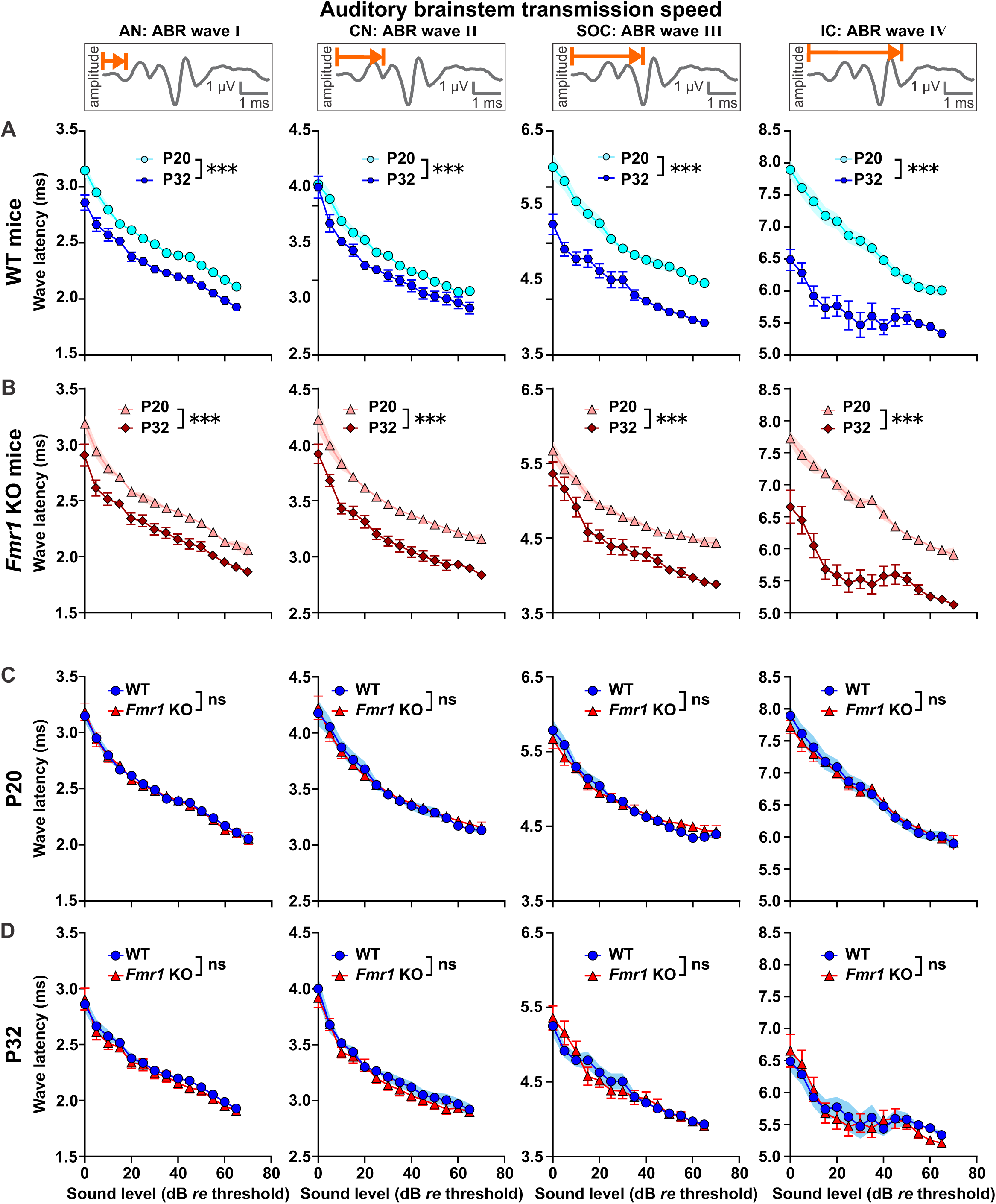
ABR wave I to IV negative peak latencies were similar in WT and *Fmr1* KO mice across development. ABR wave latencies corresponding to AN (wave I), CN (wave II), SOC (wave III), and LL and IC (wave IV) in response to pure tone stimuli (11.3 kHz) with increasing sound intensity. Note that ABR wave latencies are plotted from 0 to 65 or 0 to 70 dB *re* threshold (normalization to ABR threshold). These dB ranges were chosen to include ABRs from at least 3 males and 3 females per experimental group. ABR wave I to IV latencies decreased with age between **(A)** WT_P20 (light blue circles) and WT_P32 mice (dark blue hexagons), as well as between **(B)** *Fmr1* KO_P20 (light red triangles) and *Fmr1*_P32 mice (dark red diamonds). ABR wave I to IV latencies were similar between **(C)** WT_P20 (blue circles) and *Fmr1* KO_P20 mice (red triangles), as well as between **(D)** WT_P32 (blue circles) and *Fmr1* KO_P32 mice (red triangles). WT_P20 (*n*=16), *Fmr1* KO_P20 (*n*=15), WT_P32 (*n*=10), and *Fmr1 KO_P32* (*n*=12). Data expressed as mean (symbols) ± SEM (shaded areas or error bars). *p* values, ***p < 0.001, ns not significant.

**Table 3.** Statistical comparisons of ABR wave latencies (negative leading peaks, 0 to 65 dB re threshold) from WT mice of the following two groups: WT_P20 (*n*=16) and WT_P32 (*n*=10). This table is associated with data in Fig. 4A.

| Parameter | Source of Variation | P value | P value summary | F (DFn, DFd) | $\eta p^2$ |
| --- | --- | --- | --- | --- | --- |
| Wave I | Sex | 0.9 | ns | F(1, 21.98) = 0.01 | <0.001 |
|  | dB.re.Thr | <0.001 | *** | F(13, 280.04) = 241.84 | 0.918 |
|  | Age | <0.001 | *** | F(1, 21.98) = 33.59 | 0.604 |
|  | Sex:dB.re.Thr | 0.98 | ns | F(13, 280.05) = 0.36 | 0.017 |
|  | Sex:Age | 0.46 | ns | F(1, 21.98) = 0.55 | 0.025 |
|  | dB.re.Thr:Age | 0.66 | ns | F(13, 280.05) = 0.79 | 0.036 |
|  | Sex:dB.re.Thr:Age | 0.40 | ns | F(13, 280.05) = 1.05 | 0.047 |
| Wave II | Sex | 0.7 | ns | F(1, 21.99) = 0.15 | 0.007 |
|  | dB.re.Thr | <0.001 | *** | F(13, 280.01) = 257.54 | 0.923 |
|  | Age | <0.001 | *** | F(1, 21.99) = 17.99 | 0.450 |
|  | Sex:dB.re.Thr | 0.15 | ns | F(13, 280.01) = 1.41 | 0.062 |
|  | Sex:Age | 0.82 | ns | F(1, 21.99) = 0.05 | 0.002 |
|  | dB.re.Thr:Age | 0.05 | ns | F(13, 280.01) = 1.72 | 0.074 |
|  | Sex:dB.re.Thr:Age | 0.66 | ns | F(13, 280.01) = 0.8 | 0.036 |
| Wave III | Sex | 0.89 | ns | $F(1, 21.99) = 0.01$ | <0.001 |
| | dB.re.Thr | <0.001 | *** | $F(13, 280.03) = 210.28$ | 0.907 |
| | Age | <0.001 | *** | $F(1, 21.99) = 28.55$ | 0.565 |
| | Sex:dB.re.Thr | 0.42 | ns | $F(13, 280.03) = 1.03$ | 0.046 |
| | Sex:Age | 0.82 | ns | $F(1, 21.99) = 0.05$ | 0.002 |
| | dB.re.Thr:Age | 0.11 | ns | $F(13, 280.03) = 1.5$ | 0.065 |
| | Sex:dB.re.Thr:Age | 0.40 | ns | $F(13, 280.03) = 1.04$ | 0.046 |
| Wave IV | Sex | 0.24 | ns | $F(1, 21.99) = 1.4$ | 0.060 |
| | dB.re.Thr | <0.001 | *** | $F(13, 280.04) = 66.3$ | 0.755 |
| | Age | <0.001 | *** | $F(1, 21.99) = 57.12$ | 0.722 |
| | Sex:dB.re.Thr | 0.19 | ns | $F(13, 280.04) = 1.32$ | 0.058 |
| | Sex:Age | 0.50 | ns | $F(1, 21.99) = 0.45$ | 0.020 |
| | dB.re.Thr:Age | <0.001 | *** | $F(13, 280.04) = 9.83$ | 0.313 |
| | Sex:dB.re.Thr:Age | 0.77 | ns | $F(13, 280.04) = 0.69$ | 0.031 |

Mixed effects ART ANOVA, $p$ values, \*\*\* $p < 0.001$ , ns not significant.

**Table 4.** Statistical comparisons of ABR wave latencies (negative leading peaks, 0 to 70 dB re threshold) from *Fmr1* KO mice of the following two groups: *Fmr1* KO_P20 (*n*=15) and *Fmr1 KO_P32* (*n*=12). This table is associated with data in Fig. 4B.

| Parameter | Source of Variation | P value | P value summary | F (DFn, DFd) | $\eta p^2$ |
| --- | --- | --- | --- | --- | --- |
| Wave I | Sex | 0.22 | ns | $F(1, 23.02) = 1.52$ | 0.062 |
| | dB.re.Thr | <0.001 | *** | $F(14, 303.05) = 277.29$ | 0.928 |
| | Age | <0.001 | *** | $F(1, 23.03) = 24.5$ | 0.515 |
| | Sex:dB.re.Thr | 0.001 | ** | $F(14, 303.06) = 2.63$ | 0.108 |
| | Sex:Age | 0.77 | ns | $F(1, 23.02) = 0.08$ | 0.004 |
| | dB.re.Thr:Age | 0.09 | ns | $F(14, 303.06) = 1.54$ | 0.067 |
| | Sex:dB.re.Thr:Age | 0.83 | ns | $F(14, 303.06) = 0.64$ | 0.029 |
| Wave II | Sex | 0.06 | ns | $F(1, 23.02) = 3.69$ | 0.138 |
| | dB.re.Thr | <0.001 | *** | $F(14, 303.04) = 386.07$ | 0.947 |
| | Age | <0.001 | *** | $F(1, 23.03) = 47.99$ | 0.676 |
| | Sex:dB.re.Thr | <0.001 | *** | $F(14, 303.06) = 2.82$ | 0.115 |
| | Sex:Age | 0.45 | ns | $F(1, 23.02) = 0.57$ | 0.024 |
| | dB.re.Thr:Age | 0.83 | ns | $F(14, 303.06) = 0.63$ | 0.028 |
| | Sex:dB.re.Thr:Age | 0.90 | ns | $F(14, 303.06) = 0.54$ | 0.024 |
| Wave III | Sex | 0.76 | ns | $F(1, 23.03) = 0.09$ | 0.004 |
| | dB.re.Thr | <0.001 | *** | $F(14, 303.06) = 150.91$ | 0.875 |
| | Age | <0.001 | *** | $F(1, 23.03) = 20.64$ | 0.473 |
| | Sex:dB.re.Thr | <0.001 | *** | $F(14, 303.07) = 3.31$ | 0.133 |
| | Sex:Age | 0.33 | ns | $F(1, 23.03) = 0.95$ | 0.040 |
| | dB.re.Thr:Age | 0.34 | ns | $F(14, 303.07) = 1.11$ | 0.049 |
| | Sex:dB.re.Thr:Age | 0.74 | ns | $F(14, 303.07) = 0.72$ | 0.033 |
| Wave IV | Sex | 0.24 | ns | $F(1, 23.06) = 1.45$ | 0.059 |
| | dB.re.Thr | <0.001 | *** | $F(14, 303.18) = 75.18$ | 0.776 |
| | Age | <0.001 | *** | $F(1, 23.05) = 58.3$ | 0.717 |
| | Sex:dB.re.Thr | 0.002 | ** | $F(14, 303.15) = 2.45$ | 0.102 |
| | Sex:Age | 0.86 | ns | $F(1, 23.06) = 0.02$ | 0.001 |
| | dB.re.Thr:Age | <0.001 | *** | $F(14, 303.15) = 9.73$ | 0.310 |
| | Sex:dB.re.Thr:Age | <0.001 | *** | $F(14, 303.16) = 2.71$ | 0.112 |

Mixed effects ART ANOVA, $p$ values, \*\* $p < 0.01$ , \*\*\* $p < 0.001$ , ns not significant.

**Table 5.** *Post hoc* pairwise comparisons of ABR wave IV latencies (negative leading peaks, 0 to 70 dB re threshold) from female or male *Fmr1* KO mice of the following two groups: *Fmr1* KO_P20 (*n*=15) and *Fmr1 KO_P32* (*n*=12). This table is associated with data in Fig. 4B.

| **Contrast** | **dB** | **Estimate** | **SE** | **DF** | **t** | **Adjusted**<br>**P Value** | **P value**<br>**summary** |
| --- | --- | --- | --- | --- | --- | --- | --- |
| Female *Fmr1* KO_P20 vs<br>*Fmr1* KO_P32<br>Sex:dB.re.Thr:Age | 0 | 63.95 | 30.60 | 81.55 | 2.09 | 0.99 | ns |
|  | 5 | 103.12 | 30.60 | 81.55 | 3.37 | 0.42 | ns |
|  | 10 | 146.06 | 30.60 | 81.55 | 4.77 | 0.009 | ** |
|  | 15 | 199.63 | 30.60 | 81.55 | 6.52 | <0.001 | *** |
|  | 20 | 202.33 | 30.60 | 81.55 | 6.61 | <0.001 | *** |
|  | 25 | 189.63 | 30.60 | 81.55 | 6.20 | <0.001 | *** |
|  | 30 | 186.64 | 30.60 | 81.55 | 6.10 | <0.001 | *** |
|  | 35 | 194.11 | 30.60 | 81.55 | 6.34 | <0.001 | *** |
|  | 40 | 158.74 | 30.60 | 81.55 | 5.19 | 0.001 | ** |
|  | 45 | 120.13 | 30.60 | 81.55 | 3.93 | 0.12 | ns |
|  | 50 | 107.87 | 30.60 | 81.55 | 3.52 | 0.31 | ns |
|  | 55 | 126.89 | 30.60 | 81.55 | 4.15 | 0.06 | ns |
|  | 60 | 121.52 | 30.60 | 81.55 | 3.97 | 0.10 | ns |
|  | 65 | 122.17 | 32.15 | 96.23 | 3.80 | 0.16 | ns |
|  | 70 | 135.00 | 36.26 | 139.02 | 3.72 | 0.18 | ns |
| Male *Fmr1* KO_P20 vs<br>*Fmr1* KO_P32<br>Sex:dB.re.Thr:Age | 0 | 173.50 | 29.71 | 81.55 | 5.84 | <0.001 | *** |
|  | 5 | 140.69 | 29.71 | 81.55 | 4.74 | 0.01 | * |
|  | 10 | 176.83 | 29.71 | 81.55 | 5.95 | <0.001 | *** |
|  | 15 | 220.17 | 29.71 | 81.55 | 7.41 | <0.001 | *** |
|  | 20 | 204.75 | 29.71 | 81.55 | 6.89 | <0.001 | *** |
|  | 25 | 208.79 | 29.71 | 81.55 | 7.03 | <0.001 | *** |
|  | 30 | 165.81 | 29.71 | 81.55 | 5.58 | <0.001 | *** |
|  | 35 | 195.40 | 29.71 | 81.55 | 6.58 | <0.001 | *** |
|  | 40 | 124.88 | 29.71 | 81.55 | 4.20 | 0.05 | ns |
|  | 45 | 107.83 | 29.71 | 81.55 | 3.63 | 0.25 | ns |
|  | 50 | 100.42 | 29.71 | 81.55 | 3.38 | 0.41 | ns |
|  | 55 | 108.91 | 30.22 | 86.42 | 3.60 | 0.26 | ns |
|  | 60 | 114.17 | 30.22 | 86.42 | 3.78 | 0.17 | ns |
|  | 65 | 94.08 | 31.78 | 102.09 | 2.96 | 0.73 | ns |
|  | 70 | 112.45 | 37.10 | 159.91 | 3.03 | 0.68 | ns |

ART-C Tukey's multiple comparisons test, *p* values, \**p* < 0.05, \*\**p* < 0.01, \*\*\**p* < 0.001, ns not significant.

To assess a possible contribution of altered neuronal transmission speed on AGS susceptibility, we next compared ABR wave latencies between genotypes. In infants, response latencies were similar between WT_P20 and *Fmr1* KO_P20 mice along the ascending auditory pathway (wave I to IV, Fig. 4C, Table 6). Similarly, ABR wave latencies were not different in juvenile WT_P32 and *Fmr1* KO_P32 mice (wave I to IV, Fig. 4D, Table 7).

**Table 6.**
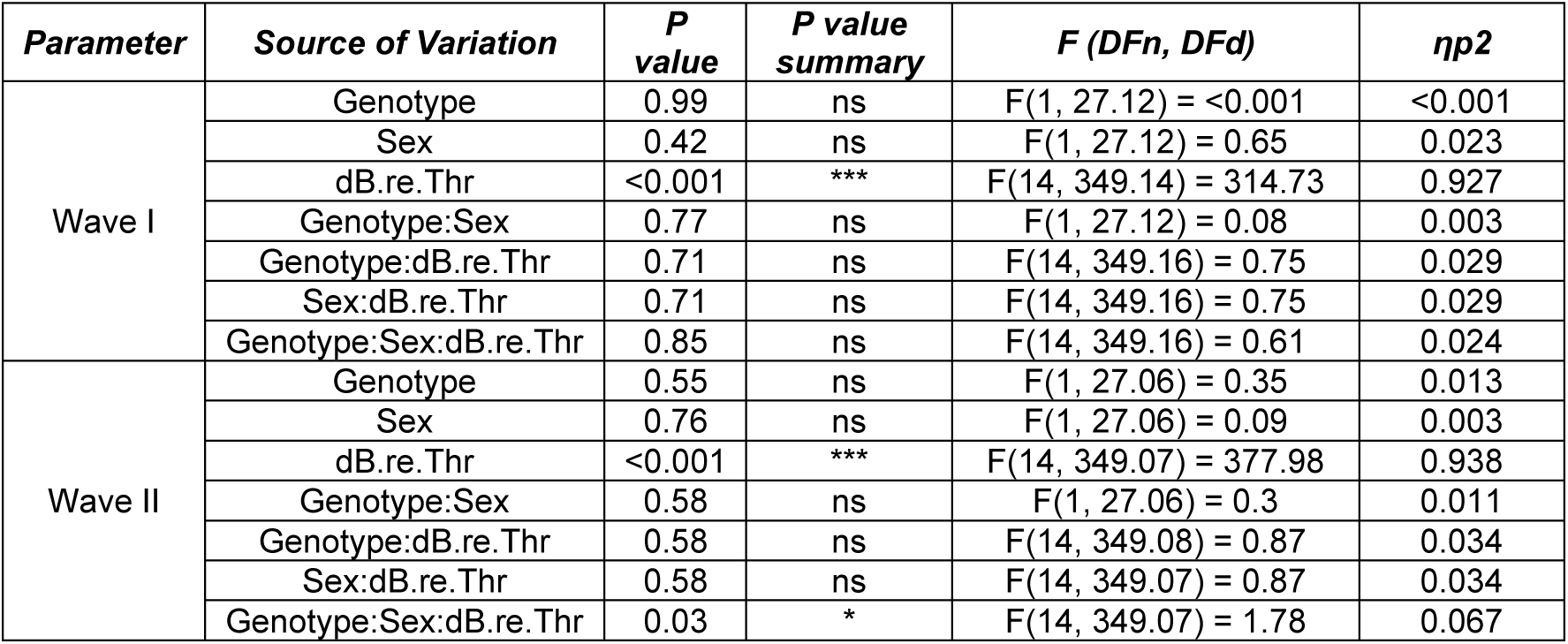
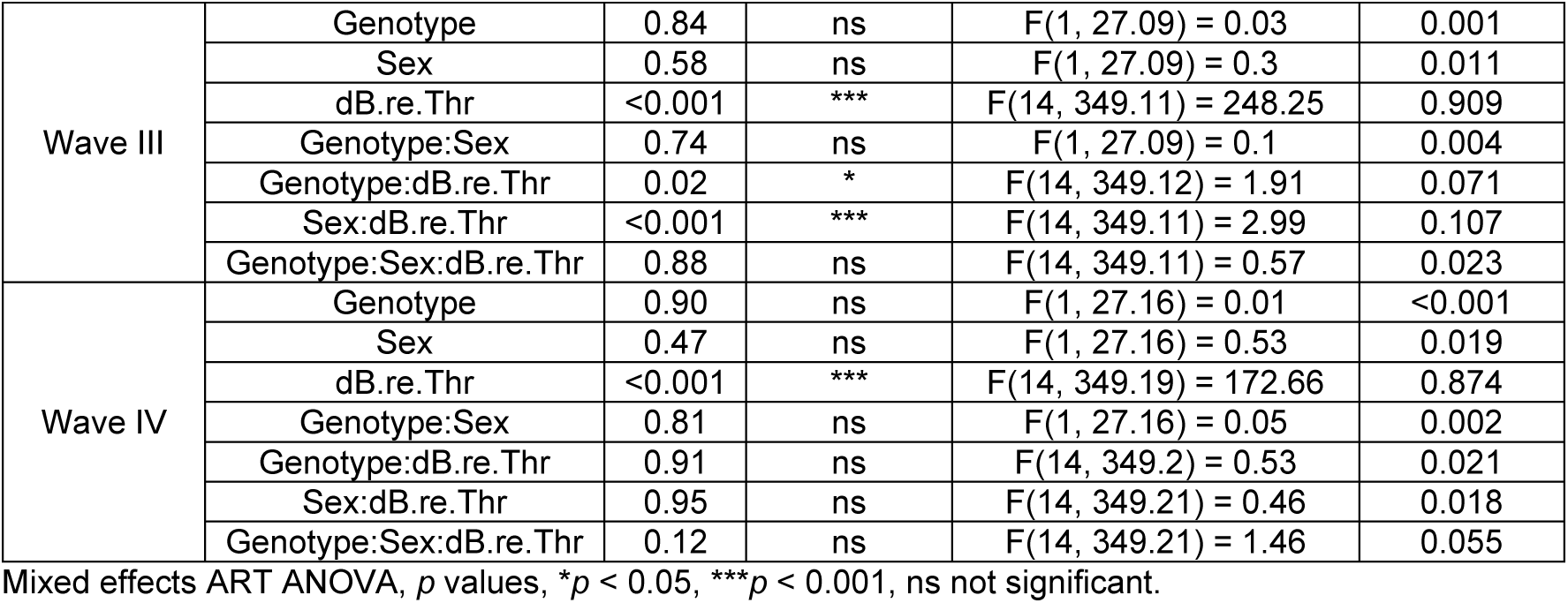
Statistical comparisons of ABR wave latencies (negative leading peaks, 0 to 70 dB re threshold) from infant mice of the following two groups: WT_P20 (*n*=16) and *Fmr1* KO_P20 (*n*=15). This table is associated with data in Fig. 4C.

**Table 7.**
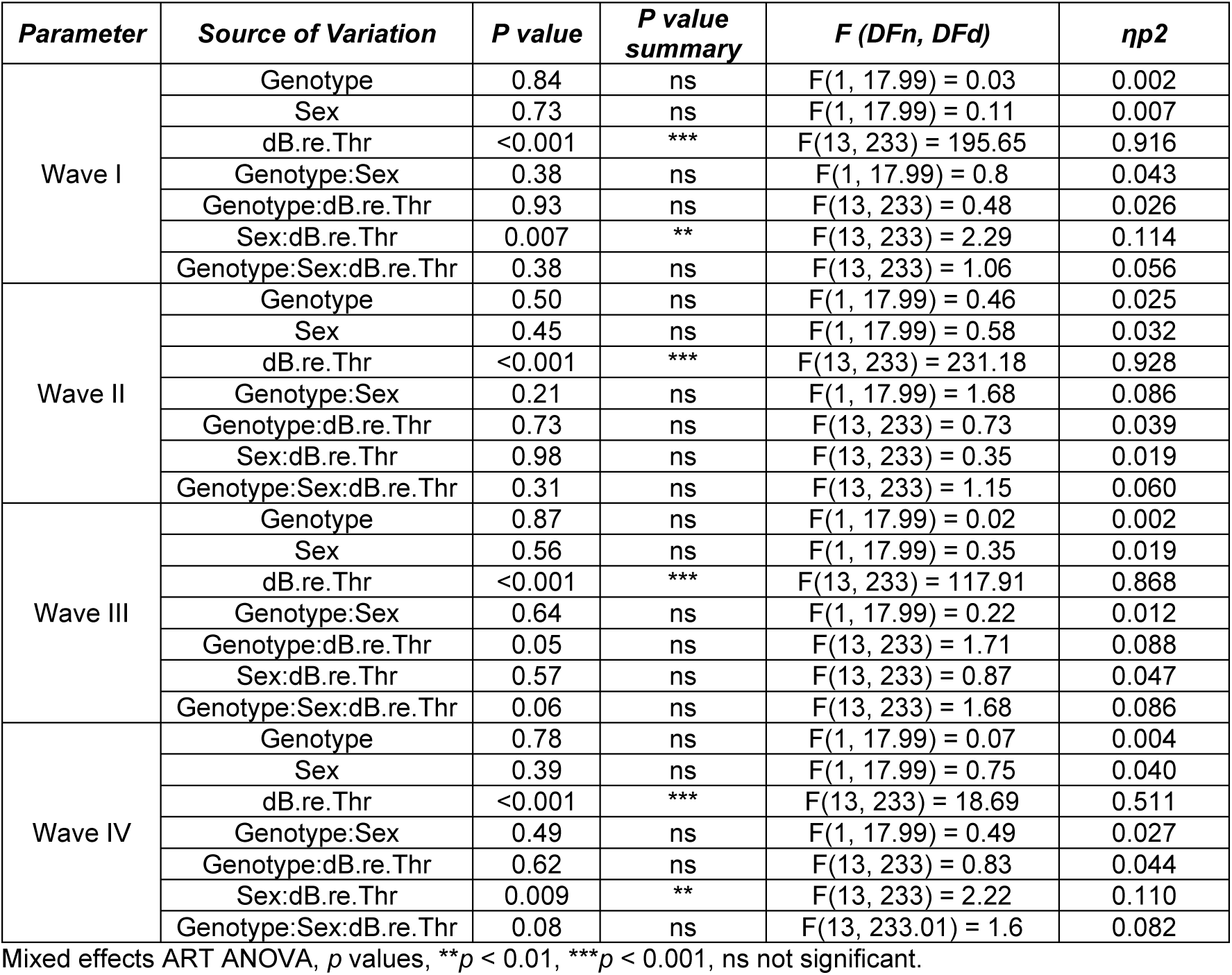
Statistical comparisons of ABR wave latencies (negative leading peaks, 0 to 65 dB re threshold) from juvenile mice of the following two groups: WT_P32 (*n*=10) and *Fmr1 KO_P32* (*n*=12). This table is associated with data in Fig. 4D.

Taken together, our results indicate that *Fmr1* mutation does not affect the maturation of neuronal transmission speed along the auditory brainstem during the observed developmental timeframe. Thus, AGS susceptibility is unlikely to be driven exclusively by alterations in neuronal transmission speed.

### 3.4. *Fmr1* deletion alters maturation of neuronal responsiveness in distinct parts of the auditory pathway

Sound-evoked ABR waveform amplitudes change proportionally to the discharge rate and the number of synchronously firing of auditory fibers (Johnson and Kiang, 1976). Therefore, changes in above-threshold ABR waveform amplitudes provide information on the neuronal responsiveness in afferent auditory fibers (Rüttiger et al., 2017). In rodents, the maturation of ABR wave amplitudes occurs non-monotonically and somewhat independent among the different waves. ABR waves I to III grow during the first ∼3-4 postnatal weeks, subsequently decline in amplitude and stabilize at adult values at ∼4-5 weeks of age; wave IV shows increased amplitude into the 4^th^ and 5^th^ postnatal week (Smith and Kraus, 1987; Song et al., 2006). To assess whether altered maturation leads to exaggerated auditory neuronal responsiveness in distinct parts of the auditory brainstem and thereby contributes to AGS susceptibility, we analyzed the amplitudes of ABR wave I to IV from *Fmr1* KO_P20, WT_P20, *Fmr1* KO_P32, and WT_P32 mice (for exemplary ABR waveforms see Supplementary Fig. 1).

Wave I amplitudes increased with age between WT_P20 and WT_P32 mice (Fig. 5A, Table 8), but not between *Fmr1* KO_P20 and *Fmr1* KO_P32 mice (Fig. 5B, Table 10). Between-genotype comparisons demonstrated that wave I amplitudes were similar in infant WT_P20 and *Fmr1* KO_P20 mice (Fig. 5C, Table 11), and amplitudes remained significantly lower in juvenile *Fmr1* KO_P32 compared with WT_P32 mice (Fig. 5D, Table 12). This indicates that the developmental increase of ABR wave I amplitudes might be delayed or absent in *Fmr1* KO mice. Furthermore, it seems unlikely that sound-evoked activity in the AN alone contributes to AGS susceptibility in these mice at the infant age.

**Fig. 5.**
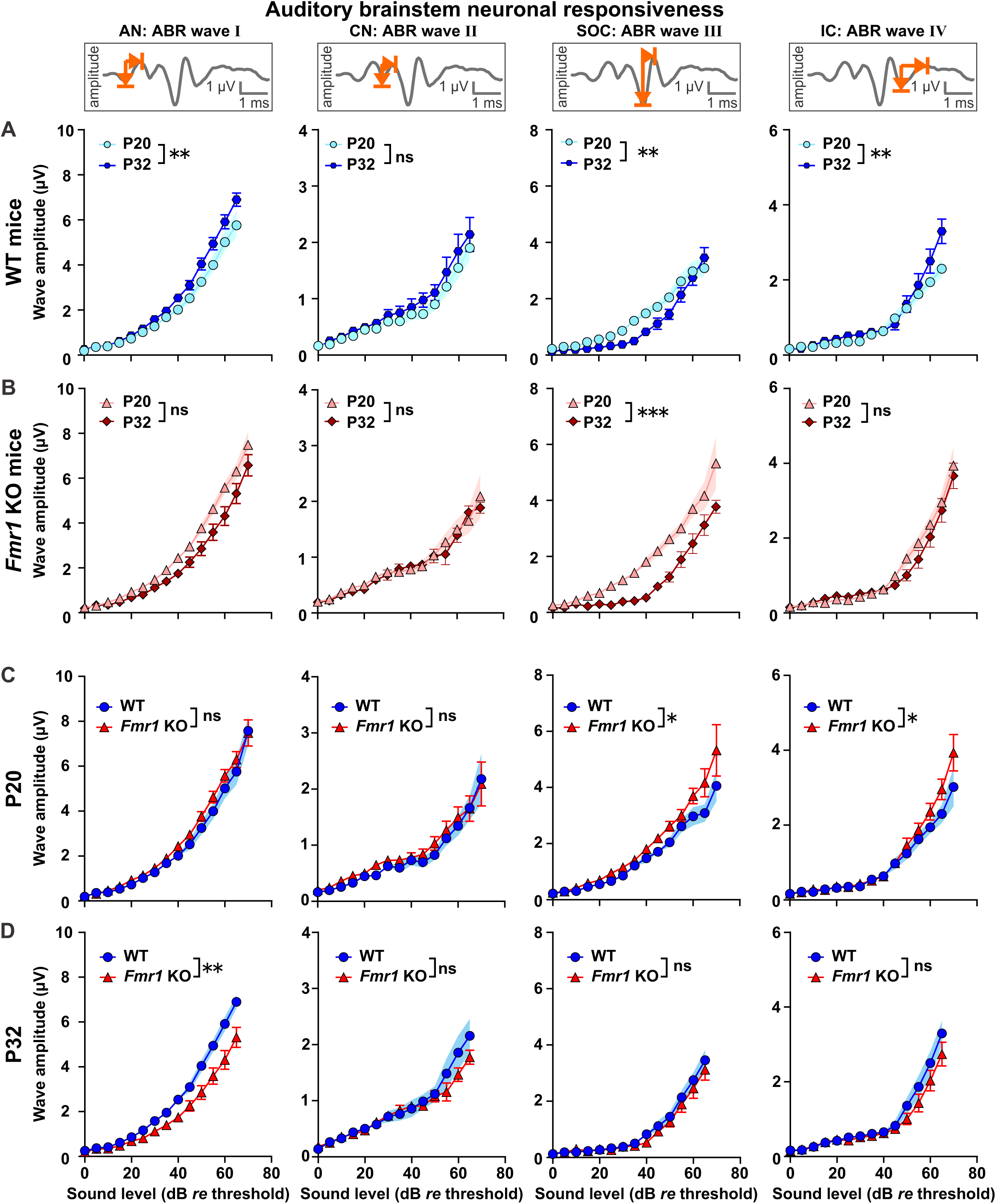
Distinct alterations in ABR wave I to IV peak-to-peak amplitudes in *Fmr1* KO mice across development. ABR wave amplitudes corresponding to AN (wave I), CN (wave II), SOC (wave III), and LL and IC (wave IV) in response to pure tone stimuli (11.3 kHz) with increasing sound intensity. Note that ABR wave amplitudes are plotted from 0 to 65 or 0 to 70 dB *re* threshold (normalization to ABR threshold). These dB ranges were chosen to include ABRs from at least 3 males and 3 females per experimental group. **(A)** During development in WTs, amplitudes of ABR wave I increased, wave II remained similar, wave III decreased, and wave IV increased in WT_P32 mice (dark blue hexagons) compared to WT_P20 (light blue circles). **(B)** During development in *Fmr1* KOs, amplitudes of ABR waves I, II, and IV remained similar, and of wave III decreased in *Fmr1*_P32 mice (dark red diamonds) compared with *Fmr1* KO_P20 (light red triangles). **(C)** Between genotypes at P20, amplitudes of ABR wave I and II were similar and of wave III and IV increased in *Fmr1* KO_P20 mice (red triangles) compared with WT_P20 (blue circles) mice. **(D)** Between genotypes at P32, amplitudes of ABR wave I were decreased and of wave II, III, and IV similar in *Fmr1* KO_P32 mice (red triangles) compared with WT_P32 (blue circles) mice. WT_P20 (*n*=16), *Fmr1* KO_P20 (*n*=15), WT_P32 (*n*=10), and *Fmr1 KO_P32* (*n*=12). Data expressed as mean (symbols) ± SEM (shaded areas or error bars). *p* values, ***p < 0.001, **p < 0.01, *p < 0.05, ns not significant.

**Table 8.**
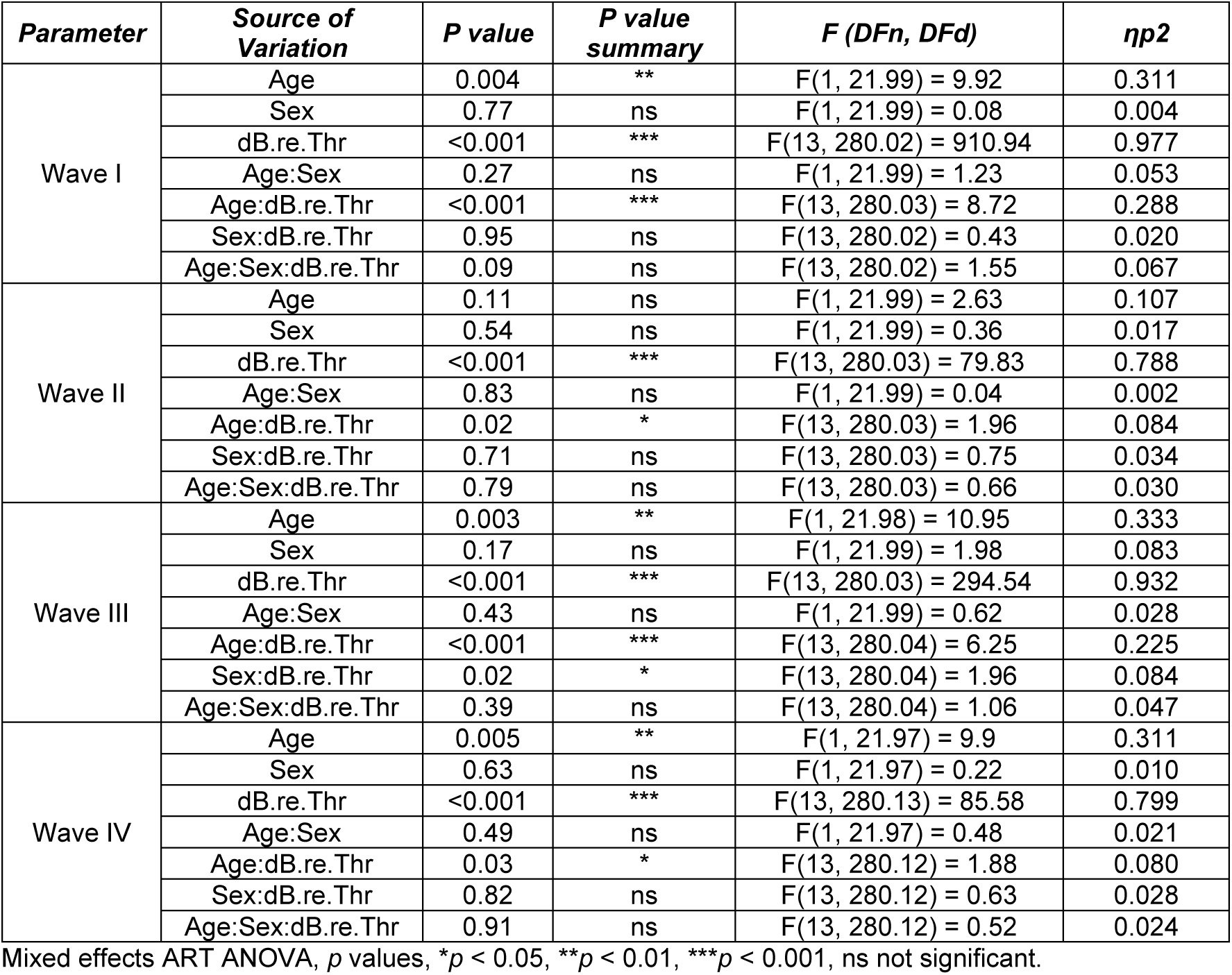
Statistical comparisons of ABR wave amplitudes (peak-to-peak, 0 to 65 dB re threshold) from WT mice of the following two groups: WT_P20 (*n*=16) and WT_P32 (*n*=10). This table is associated with data in Fig. 5A.

**Table 9.**
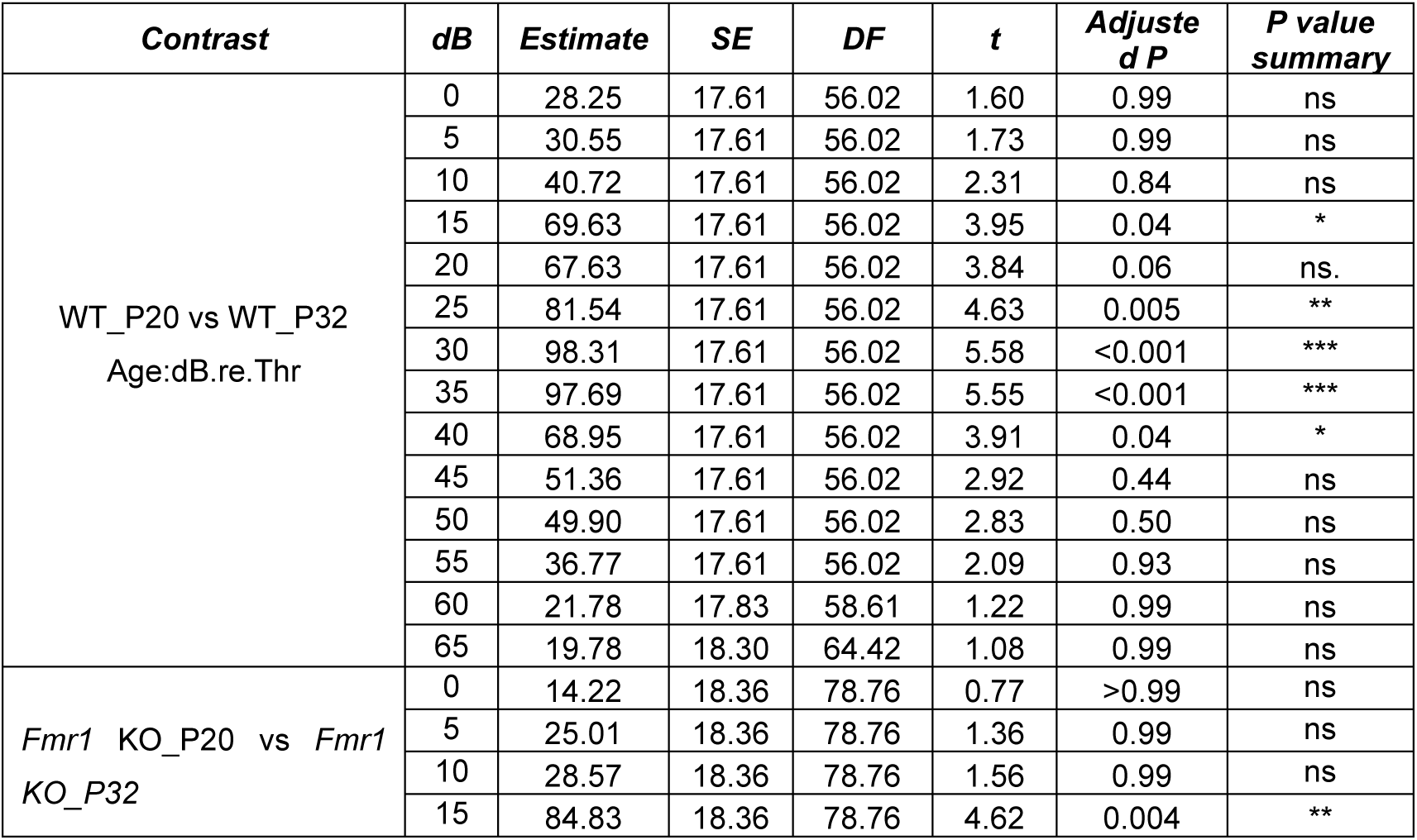
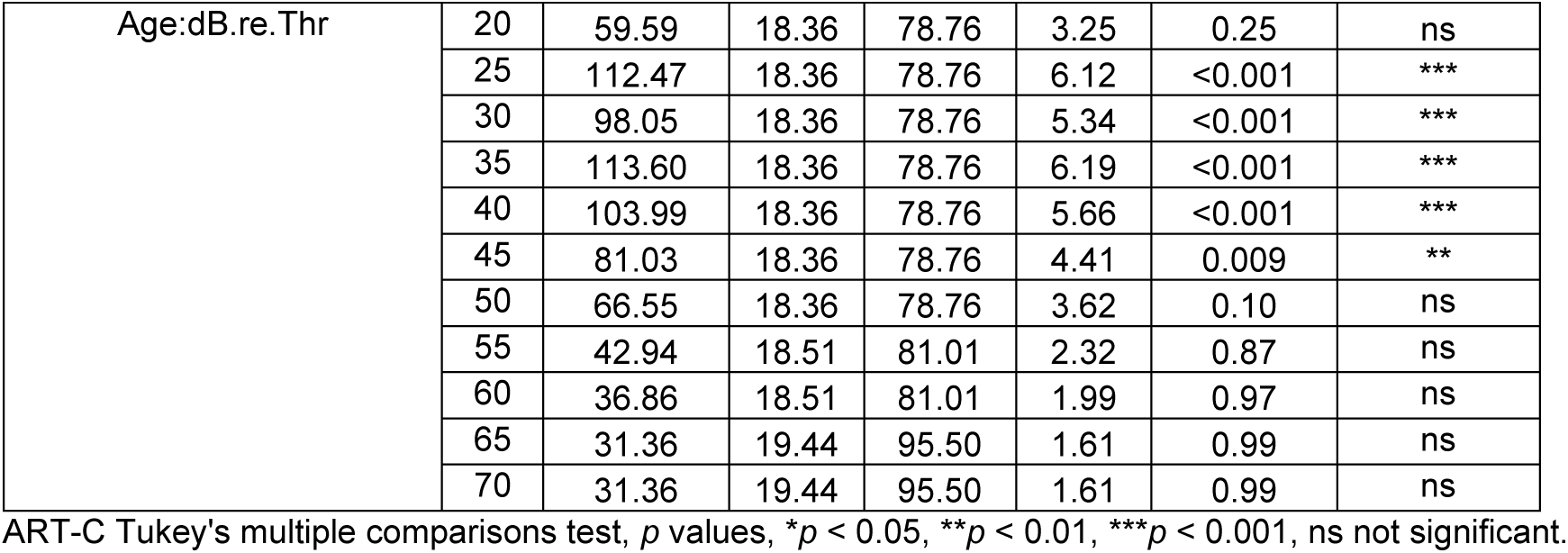
*Post hoc* pairwise comparisons of ABR wave III amplitudes (peak-to-peak) for contrast age x dB re threshold from mice of the following groups: WT_P20 (*n*=16) and WT_P32 (*n*=10) or *Fmr1* KO_P20 (*n*=15) and *Fmr1 KO_P32* (*n*=12). This table is associated with data in Fig. 5A.

**Table 10.** Statistical comparisons of ABR wave amplitudes (peak-to-peak, 0 to 70 dB re threshold) from *Fmr1* KO mice of the following two groups: *Fmr1* KO_P20 (*n*=15) and *Fmr1 KO_P32* (*n*=12). This table is associated with data in Fig. 5B.

| Parameter | Source of Variation | P value | P value summary | F (DFn, DFd) | $\eta^2$ |
| --- | --- | --- | --- | --- | --- |
| Wave I | Age | 0.27 | ns | F(1, 23.03) = 1.23 | 0.051 |
|  | Sex | 0.57 | ns | F(1, 23.03) = 0.31 | 0.014 |
|  | dB.re.Thr | <0.001 | *** | F(14, 303.06) = 1158.05 | 0.982 |
|  | Age:Sex | 0.18 | ns | F(1, 23.03) = 1.85 | 0.075 |
|  | Age:dB.re.Thr | <0.001 | *** | F(14, 303.07) = 4.02 | 0.157 |
|  | Sex:dB.re.Thr | 0.83 | ns | F(14, 303.07) = 0.63 | 0.028 |
|  | Age:Sex:dB.re.Thr | 0.27 | ** | F(14, 303.07) = 2.67 | 0.110 |
| Wave II | Age | 0.84 | ns | F(1, 23.08) = 0.04 | 0.002 |
|  | Sex | 0.69 | ns | F(1, 23.08) = 0.15 | 0.007 |
|  | dB.re.Thr | <0.001 | *** | F(14, 303.18) = 92.73 | 0.811 |
|  | Age:Sex | 0.23 | ns | F(1, 23.08) = 1.49 | 0.061 |
|  | Age:dB.re.Thr | 0.94 | ns | F(14, 303.2) = 0.46 | 0.021 |
|  | Sex:dB.re.Thr | 0.85 | ns | F(14, 303.2) = 0.61 | 0.028 |
|  | Age:Sex:dB.re.Thr | 0.03 | * | F(14, 303.2) = 1.85 | 0.079 |
| Wave III | Age | <0.001 | *** | F(1, 23.07) = 17.98 | 0.438 |
|  | Sex | 0.09 | ns | F(1, 23.06) = 3.09 | 0.118 |
|  | dB.re.Thr | <0.001 | *** | F(14, 303.12) = 261.59 | 0.924 |
|  | Age:Sex | 0.33 | ns | F(1, 23.06) = 0.96 | 0.040 |
|  | Age:dB.re.Thr | <0.001 | *** | F(14, 303.15) = 9.65 | 0.308 |
|  | Sex:dB.re.Thr | <0.001 | *** | F(14, 303.14) = 4.12 | 0.160 |
|  | Age:Sex:dB.re.Thr | 0.47 | ns | F(14, 303.14) = 0.97 | 0.043 |
| Wave IV | Age | 0.60 | ns | F(1, 23.09) = 0.26 | 0.012 |
|  | Sex | 0.34 | ns | F(1, 23.09) = 0.94 | 0.039 |
|  | dB.re.Thr | <0.001 | *** | F(14, 303.32) = 95.4 | 0.815 |
|  | Age:Sex | 0.88 | ns | F(1, 23.09) = 0.02 | 0.001 |
|  | Age:dB.re.Thr | 0.09 | ns | F(14, 303.21) = 1.53 | 0.066 |
|  | Sex:dB.re.Thr | 0.06 | ns | F(14, 303.24) = 1.63 | 0.070 |
|  | Age:Sex:dB.re.Thr | 0.47 | ns | F(14, 303.23) = 0.97 | 0.043 |

Mixed effects ART ANOVA, $p$ values, $p < 0.05$ , \*\* $p < 0.01$ , \*\*\* $p < 0.001$ , ns not significant.

**Table 11.**
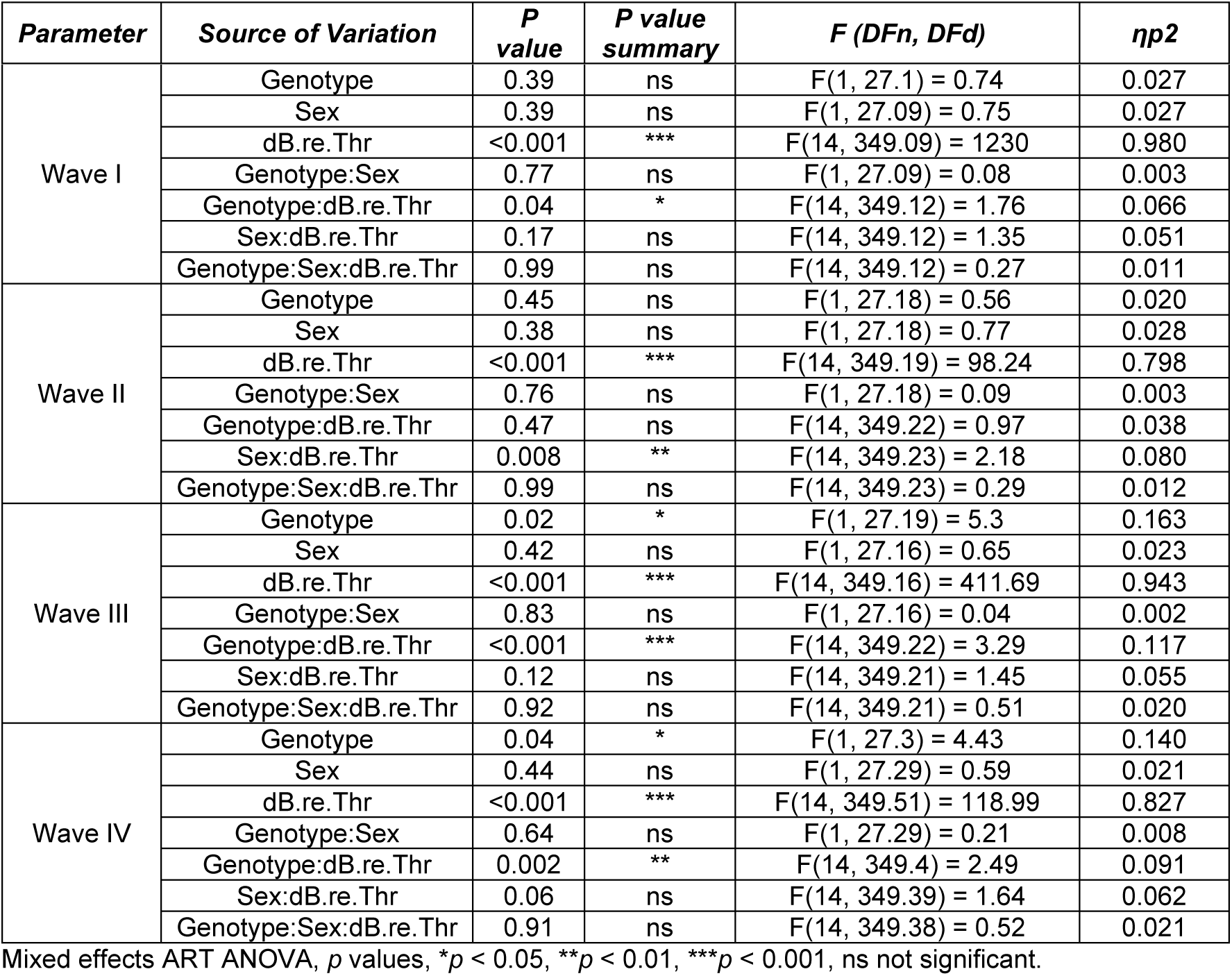
Statistical comparisons of ABR wave amplitudes (peak-to-peak, 0 to 70 dB re threshold) from infant mice of the following two groups: WT_P20 (*n*=16) and *Fmr1* KO_P20 (*n*=15). This table is associated with data in Fig. 5C.

**Table 12.**
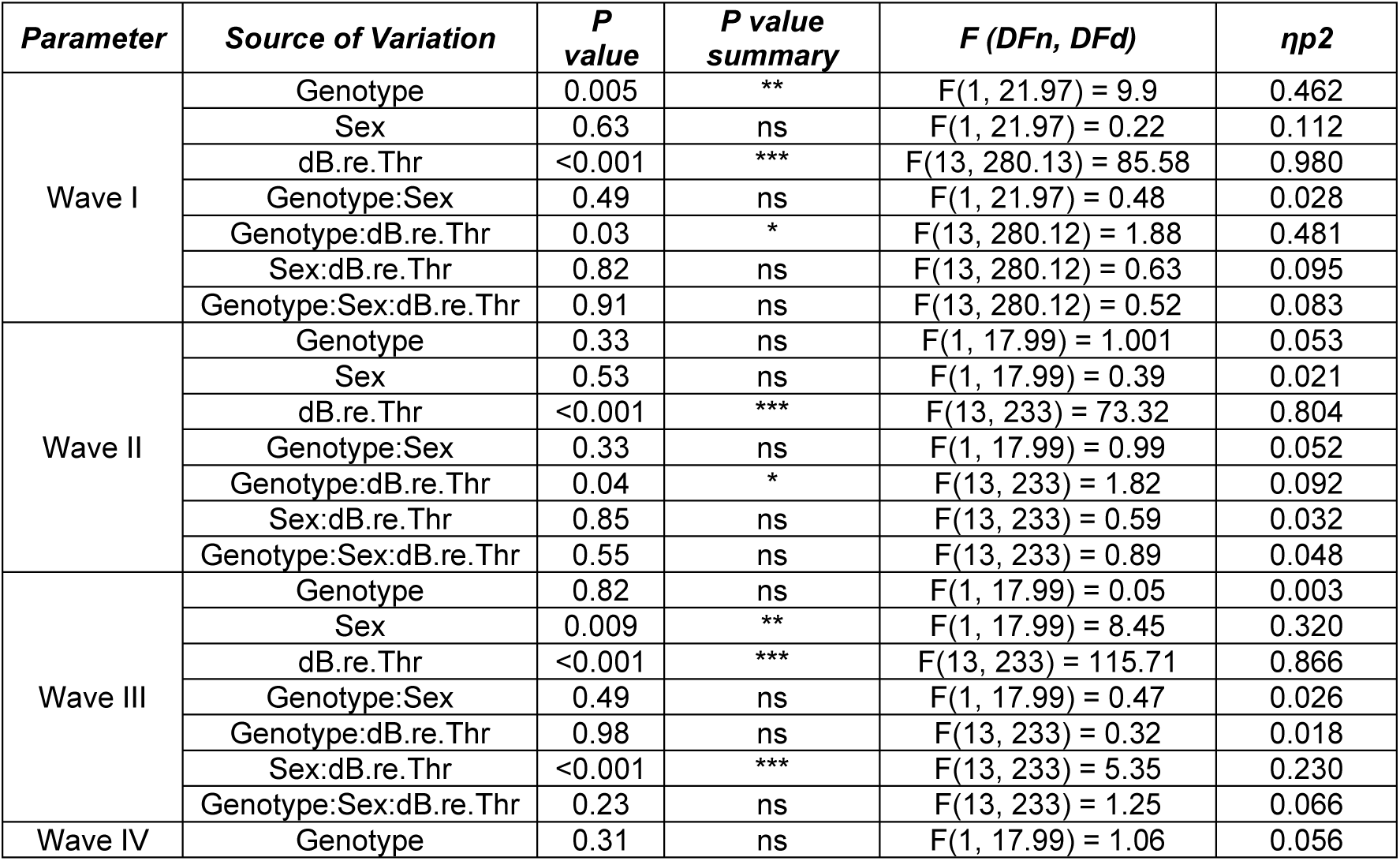
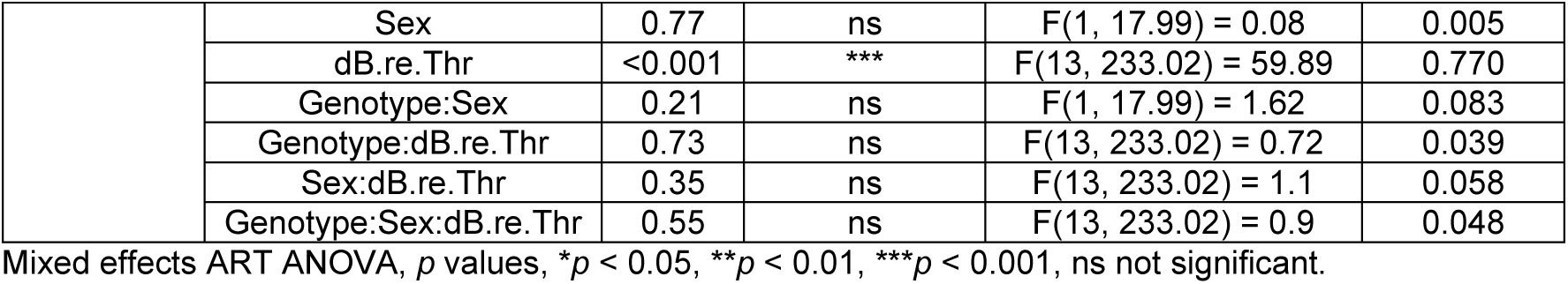
Statistical comparisons of ABR wave amplitudes (peak-to-peak, 0 to 65 dB re threshold) from juvenile mice of the following two groups: WT_P32 (*n*=10) and *Fmr1 KO_P32* (*n*=12). This table is associated with data in Fig. 5D.

Wave II amplitudes remained similar between WT_P20 and WT_P32 mice (Fig. 5A, Table 8), or *Fmr1* KO_P20 and *Fmr1* KO_P32 mice (Fig. 5B, Table 10). Amplitudes were also not different in infant WT_P20 and *Fmr1* KO_P20 mice (Fig. 5C, Table 11), or juvenile *Fmr1* KO_P32 and WT_P32 mice (Fig. 5D, Table 12). Taken together, it appears that *Fmr1* mutation did not alter wave II amplitude maturation in the observed developmental timeframe. Therefore, summed sound-evoked activity at the level of the CN likely plays only a minor role in AGS expression.

For ABR wave III, there was a reduction in amplitudes between WT_P20 and WT_P32 mice (Fig. 5A, Table 8), as well as between *Fmr1* KO_P20 and *Fmr1* KO_P32 mice (Fig. 5B, Table 10). In both WT and *Fmr1* KO mice, this developmental decrease of wave III amplitudes occurred predominantly within sound levels of ∼15 to 45 dB *re* threshold (Table 9). On average, this amplitude reduction was more pronounced in *Fmr1* KO (mean±SEM, 0.79±0.07 μV) compared with WT mice (0.51±0.08 μV, unpaired *t* test, *p* = 0.013, *t*=2.72, df=20, mean difference −0.28, 95% confidence interval −0.5021 to −0.06692). Infant *Fmr1* KO_P20 mice showed significantly elevated wave III amplitudes compared to WT_P20 mice (Fig. 5C, Table 11). By P32, amplitude values in *Fmr1* KO mice had stabilized at WT levels (Fig. 5D, Table 12). Different from other ABR parameters, there was a difference in sex for ABR wave III amplitudes at P32, with males showing higher amplitudes than females, independent of genotype (Table 12). The literature on sex differences in ABR wave amplitudes appears inconsistent (Guimaraes et al., 2004; Hunter and Willott, 1987; Kim et al., 2023; Lozier et al., 2023), emphasizing the need to incorporate this biological variable in auditory research to improve the accuracy and generalizability of results. In summary, these findings indicate that *Fmr1* mutation leads to delayed developmental decrease, *i.e*., maturation, of wave III amplitudes at the infant age. This increased neuronal responsiveness at the level of the SOC during early auditory development might contribute to AGS susceptibility.

ABR wave IV showed increased amplitudes in WT_P32 compared with WT_P20 mice (Fig. 5A, Table 8), but no difference between *Fmr1* KO_P32 and *Fmr1* KO_P20 mice (Fig. 5B, Table 10). In infancy, *Fmr1* KO_P20 mice had significantly elevated wave IV amplitudes compared with WT_P20 mice (Fig. 5C, Table 11). In *Fmr1* KO_P32 mice, amplitude values were similar to WT_P32 levels (Fig. 5D, Table 12). This suggests that *Fmr1* mutation disturbs the timely maturation of wave IV amplitudes, resulting in a premature increase in the response during early auditory development. This exaggerated sound-evoked activity at the level of the LL and IC might contribute to the AGS expression.

To further assess the contribution of increased neuronal responsiveness across the ascending auditory pathway to AGS, we next pooled the ABR wave amplitudes of infants into three groups based on AGS phenotypes, *i.e*., WT_P20 (all with no response), *Fmr1* KO_P20 with no response (AGS1), or *Fmr1* KO_P20 mice with any response (AGS2-4, including wild running, seizure, and respiratory arrest). We extracted the dynamic range of wave level-amplitude functions (dB range of rapid growth) for individual animals based on the 99% confidence intervals. We then compared the slopes of ABR wave I to IV amplitude growth functions using linear regression as a measure for the efficiency and sensitivity of the neural response to increasing sound intensity. The slopes of either ABR wave I and II growth functions were not significantly different between the three groups (Fig. 6 A, wave I, WT_P20: Y = 0.1229*X - 2.324, *Fmr1* KO_P20 AGS1: Y = 0.1452*X - 3.317, *Fmr1* KO_P20 AGS2-4: Y = 0.1231*X - 1.901, F = 1.023, DFn = 2, DFd = 26, *p* = 0.37; wave II, WT_P20: Y = 0.02377*X + 0.09261, *Fmr1* KO_P20 AGS1: Y = 0.02728*X + 0.05241, *Fmr1* KO_P20 AGS2-4: Y = 0.02464*X + 0.08063, F = 0.2705, DFn = 2, DFd = 37, *p* = 0.76). Surprisingly, the slopes of ABR wave III were also not significantly different, though there was a slight increase in steepness with AGS susceptibility (Fig. 6A, WT_P20: Y = 0.06513*X - 0.7204, *Fmr1* KO_P20 AGS1: Y = 0.07350*X - 0.4503, *Fmr1* KO_P20 AGS2-4: Y = 0.08416*X - 1.008, F = 1.331, DFn = 2, DFd = 31, *p* = 0.27). In contrast to waves I to III, the slopes of wave IV amplitude growth functions in *Fmr1* KO_P20 mice with any AGS response were significantly steeper than in WT_P20 mice (Fig. 6A, WT_P20: Y = 0.04939*X - 0.9198, *Fmr1* KO_P20 AGS2-4: Y = 0.08171*X - 2.139, F = 6.408, DFn = 1, DFd = 16, *p* = 0.022), whereas those in *Fmr1* KO_P20 mice with no response in the AGS test were not different (Y = 0.05520*X - 0.7538, F = 0.1690, DFn = 1, DFd = 14, *p* = 0.68). This indicates that the auditory midbrain at the level of the LL and IC in mice with AGS responded more intensely or showed a more rapid increase in response amplitude to changes in sound level compared to WT mice, which did not exhibit any AGS response.

**Fig. 6.**
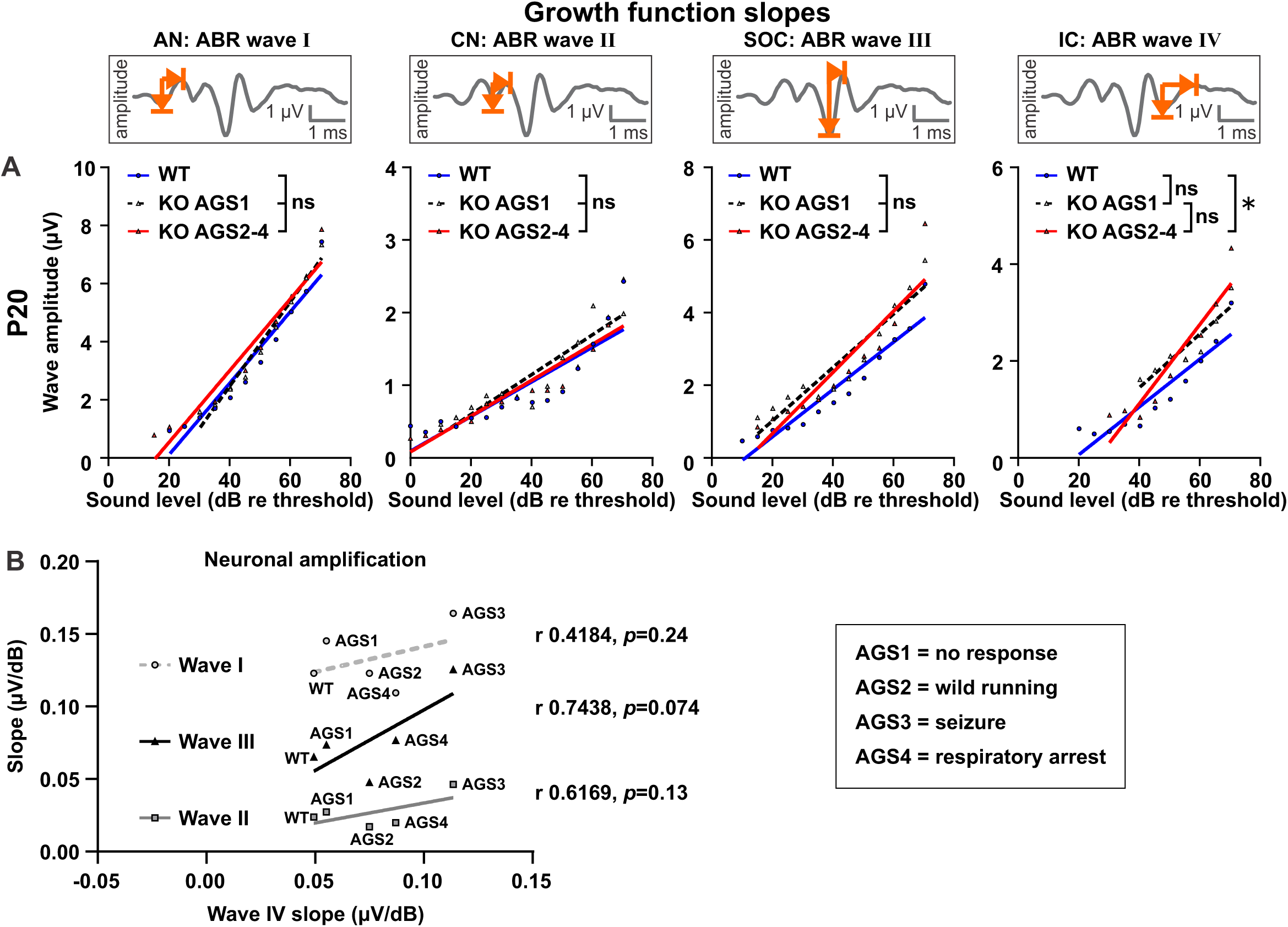
Slope analysis of ABR wave amplitude growth functions in P20 mice depending on AGS susceptibility. **(A)** The dynamic ranges of ABR wave I, II, III, and IV level-amplitude functions (dB range of rapid growth) were extracted for individual animals based on the 99% confidence intervals and pooled for mice depending on AGS phenotype. Slopes of ABR wave growth function were compared between WT_P20 controls (blue line), *Fmr1* KO_P20 AGS1 (no response; black dashed line), and *Fmr1* KO_P20 AGS2-4 mice (wild running, seizure, respiratory arrest; red line) using linear regression. Slopes were not significantly different between the three groups for ABR wave I (WT_P20: Y = 0.1229*X - 2.324, *Fmr1* KO_P20 AGS1: Y = 0.1452*X - 3.317, *Fmr1* KO_P20 AGS2-4: Y = 0.1231*X - 1.901, F = 1.023, DFn = 2, DFd = 26, *p* = 0.37), wave II (WT_P20: Y = 0.02377*X + 0.09261, *Fmr1* KO_P20 AGS1: Y = 0.02728*X + 0.05241, *Fmr1* KO_P20 AGS2-4: Y = 0.02464*X + 0.08063, F = 0.2705, DFn = 2, DFd = 37, *p* = 0.76), and wave III (WT_P20: Y = 0.06513*X - 0.7204, *Fmr1* KO_P20 AGS1: Y = 0.07350*X - 0.4503, *Fmr1* KO_P20 AGS2-4: Y = 0.08416*X - 1.008, F = 1.331, DFn = 2, DFd = 31, *p* = 0.27). Slopes of ABR wave IV amplitude growth functions were significantly different (WT_P20: Y = 0.04939*X - 0.9198, *Fmr1* KO_P20 AGS21 Y = 0.05520*X - 0.7538, *Fmr1* KO_P20 AGS2-4: Y = 0.08171*X - 2.139, F = 3.702, DFn = 2, DFd = 21, *p* = 0.04), in particular between WT_P20 and *Fmr1* KO_P20 AGS2-4 mice (F = 6.408, DFn = 1, DFd = 16, *p* = 0.022). Slopes between WT_P20 and *Fmr1* KO_P20 AGS1 (F = 0.1690, DFn = 1, DFd = 14, *p* = 0.68) as well as *Fmr1* KO_P20 AGS1 and *Fmr1* KO_P20 AGS2-4 mice (F = 2.278, DFn = 1, DFd = 12, *p* = 0.15) were not different. **(B)** Slopes of ABR wave IV correlated with slopes of wave I (gray circles and dashed line), II (gray squares and line), or III (black triangles and line) amplitude growth functions from WT_P20, *Fmr1* KO_P20 AGS1 (no response), AGS2 (wild running), AGS3 (seizure), and AGS4 (respiratory arrest) mice. Slopes of wave I and wave II showed no statistically significant correlation with wave IV (wave I v IV: Pearson r 0.4184, one-tailed *p* = 0.24; II v IV: Pearson r 0.6169, one-tailed *p* = 0.13). Slopes of wave III and IV showed a strong positive correlation that reached a statistical trend (wave III v IV: Pearson r 0.7438, one-tailed *p* = 0.074). **(A)** WT_P20 (*n*=16), *Fmr1* KO_P20 AGS2-4 (*n*=11), **(B)** WT_P20 (*n*=16), *Fmr1* KO_P20 AGS1 (*n*=4), KO_P20 AGS2 (*n*=3), KO_P20 AGS3 (*n*=3), KO_P20 AGS4 (*n*=5). Data expressed as mean (symbols) and linear regression lines. Regression lines in **(B)** are only for visual clarity. *p* values, *p < 0.05, ns not significant.

We next aimed to determine if this hyperresponsiveness originated in the midbrain, or if it may be relayed from the lower auditory structures. To this end, we correlated the slopes of ABR wave I, II, or III amplitude growth functions with those of wave IV from WT_P20, *Fmr1* KO_P20 AGS1, 2, 3, and 4 mice (Fig. 6B). We assumed that if there was no or weak correlation, then the hyperresponsiveness originated in the midbrain. If there was a strong positive correlation, then the hyperresponsiveness may be relayed from lower auditory structures. Neither the slope of wave I nor the slope of wave II showed a statistically significant correlation with the slope of wave IV (Fig 6B, wave I v IV: Pearson r 0.4184, one-tailed *p* = 0.24; II v IV: Pearson r 0.6169, one-tailed *p* = 0.13). The strong correlation between the slopes of waves III and IV reached a statistical trend (Fig. 6B, wave III v IV: Pearson r 0.7438, one-tailed *p* = 0.074). These results suggest that auditory hyperresponsiveness underlying AGS may, at least in part, originate in the lower auditory brainstem and then be relayed to and pathologically amplified in the auditory midbrain.

Taken together, these results indicate that *Fmr1* plays an important role in the proper maturation of neuronal responsiveness in distinct parts of the ascending auditory pathway, particularly at the level of the AN, SOC, as well as the LL and IC. Among these, increased responsiveness in the SOC as well as LL and IC at a time that coincides with early auditory development may underlie susceptibility to AGS and contribute to sensory hypersensitivity in FXS.

### 3.5. Increased number of cFos positive cells in the IC of in *Fmr1* KO mice

cFos expression has been widely used as a reliable histological marker to map neuronal activity in the central nervous system, including stimulus-related responses on a single-cell level in the auditory pathway (Ehret and Fischer, 1991; Moreno-Paublete et al., 2017; Yang et al., 2005). Previous studies have suggested that an increase in sound-evoked cFos-positive cell density in specific auditory nuclei might be a correlate of sensory hypersensitivity in early development (Chen and Toth, 2001; Nguyen et al., 2020; Yang et al., 2020). We stained coronal brain sections containing the IC from a separate cohort of infant and juvenile WT and *Fmr1* KO mice to determine if hyperresponsiveness in the auditory midbrain during early development is associated with increased cFos expression. Before brain extraction, the animals from both genotypes and age groups were sound-exposed to the same siren as in the AGS test (Fig. 1E). Microscopic images of the central nucleus of the IC (Fig. 7A), in an area assumed to represent a ∼8-40 kHz sound frequency region (Ryugo and Milinkeviciute, 2023), were collected and analyzed for cFos immunoreactive labelling (Fig. 7C) in a semi-automated fashion using the ImageJ/Fiji tool ‘Quanty-cFos’ (Beretta et al., 2023). cFos-positive cell counts from WT_P20, *Fmr1* KO_P20, WT_P32, and *Fmr1* KO_P32 mice showed a genotype effect (Fig. 7B, Table 13), with a higher cFos-positive cell density in *Fmr1* KO mice (median ± IQR, 77.5 ± 56.0-99.2) compared with WT mice (40.3 ± 25.0-66.8). *Post hoc* comparisons showed that this difference was particularly pronounced between *Fmr1* KO_P20 (77.5 ± 70.5-232.0) and WT_P20 mice (27.5 ± 15.9-61.1; Fig. 7B, Table 14). In contrast to infant mice, the difference was not statistically significant in juvenile *Fmr1* KO_P32 (79.5 ± 38.9-85.8) and WT_P32 mice (53.5, ± 37.1-84.4). Though the interaction between age x genotype of cFos-positive cell counts was not significant (Fig. 7B, Table 13), it is worth noting that the direction of age-related changes in cFos labelling matched those of ABR wave IV amplitudes in either WT and *Fmr1* KO mice: between P20 and P32 both cFos-positive cell density and ABR wave IV amplitudes increased in WT mice (Fig. 5A, 7B), whereas both remained similar in *Fmr1* KO mice (Fig. 5B, 7B). Interestingly, we did not observe a systematic pattern in cFos expression in *Fmr1* KO_P20 mice, regardless of whether they showed no behavioral response in the AGS test or exhibited any response (here including wild running and seizure, Fig. 7B). However, this observation should be interpreted with caution due to the small sample size.

**Fig. 7.**
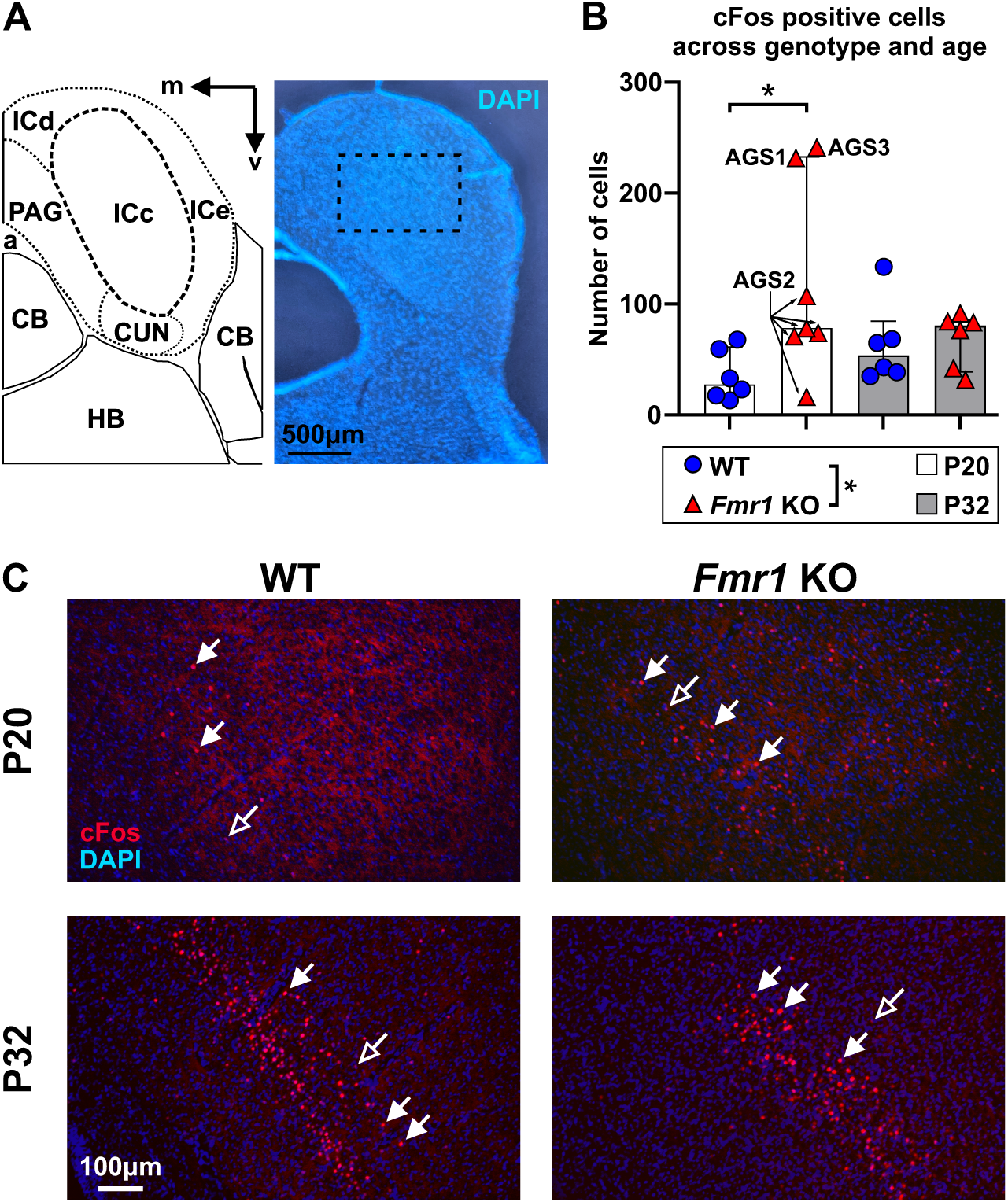
Number of cFos positive cells in the IC was increased in infant *Fmr1* KO mice. **(A)** Schematic overview and coronal section of a mouse brain showing the IC. Cell nuclei were stained with DAPI (blue), and the dashed rectangle indicates the region where the IC was imaged at higher magnification for cFos quantification. Please note that the cerebellum was incomplete in the coronal section. Scale bar, 500 μm. Abbreviations: a, cerebral aqueduct; CB, cerebellum; CUN, cuneiform nucleus; HB, hindbrain; ICc, inferior colliculus, central nucleus; ICd, inferior colliculus, dorsal nucleus; ICe, inferior colliculus, external nucleus; m, medial; PAG, periaqueductal gray; v, ventral. **(B)** Quantification of cFos immunopositive cells in the IC of infant (white bars) or juvenile (gray bars) WT (blue circles) and *Fmr1* KO (red triangles) mice. The number of cFos positive cells was significantly different between genotypes. In particular, the cFos positive cell density was higher in *Fmr1* KO_P20 compared with WT_P20 mice. **(C)** Representative coronal brain sections containing the IC stained with antibodies against cFos (red). Immunopositive dots were counted to estimate the number of cFos positive cells in the ICc. The effect of *Fmr1* mutation on the number of cFos positive cell counts were analyzed in infant (P20, upper panel) and juvenile (P32, lower panel) WT (left) and *Fmr1* KO mice (right) after exposure to the same sound used in the AGS test. Filled arrows indicate cFos positive cells and open arrows indicate cFos negative cells. Cell nuclei were stained with DAPI (blue). Scale bars, **(A)** 500 μm, **(C)** 100 μm. AGS1 = no response, AGS2 = wild running, AGS 3= seizure. WT_P20 (*n*=6), *Fmr1* KO_P20 (*n*=7), WT_P32 (*n*=6), and *Fmr1 KO_P32* (*n*=6). Data expressed as median (bars) ± IQR (error bars) and individual animals (symbols). *p* value, * *p* < 0.05.

**Table 13.** Statistical comparisons of number of cFos positive cells from mice of the following four groups: WT_P20 (*n*=6), *Fmr1* KO_P20 (*n*=7), WT_P32 (*n*=6), and *Fmr1 KO_P32* (*n*=6). This table is associated with data in Fig. 7B.

| Parameter | Source of Variation | P value | P value summary | F (DFn, DFd) | $\eta^2$ |
| --- | --- | --- | --- | --- | --- |
| cFos positive cells | Sex | 0.77 | ns | $F(1, 17) = 0.08$ | 0.005 |
| | Genotype | 0.04 | * | $F(1, 17) = 4.45$ | 0.208 |
| | Age | 0.94 | ns | $F(1, 17) = 0.004$ | >0.001 |
| | Sex:Genotype | 0.75 | ns | $F(1, 17) = 0.1$ | 0.006 |
| | Sex:Age | 0.86 | ns | $F(1, 17) = 0.02$ | 0.002 |
| | Genotype:Age | 0.11 | ns | $F(1, 17) = 2.78$ | 0.141 |
| | Sex:Genotype:Age | 0.59 | ns | $F(1, 17) = 0.29$ | 0.017 |

Three-way ART ANOVA, $p$ values, \* $p < 0.05$ , ns not significant.

**Table 14.**
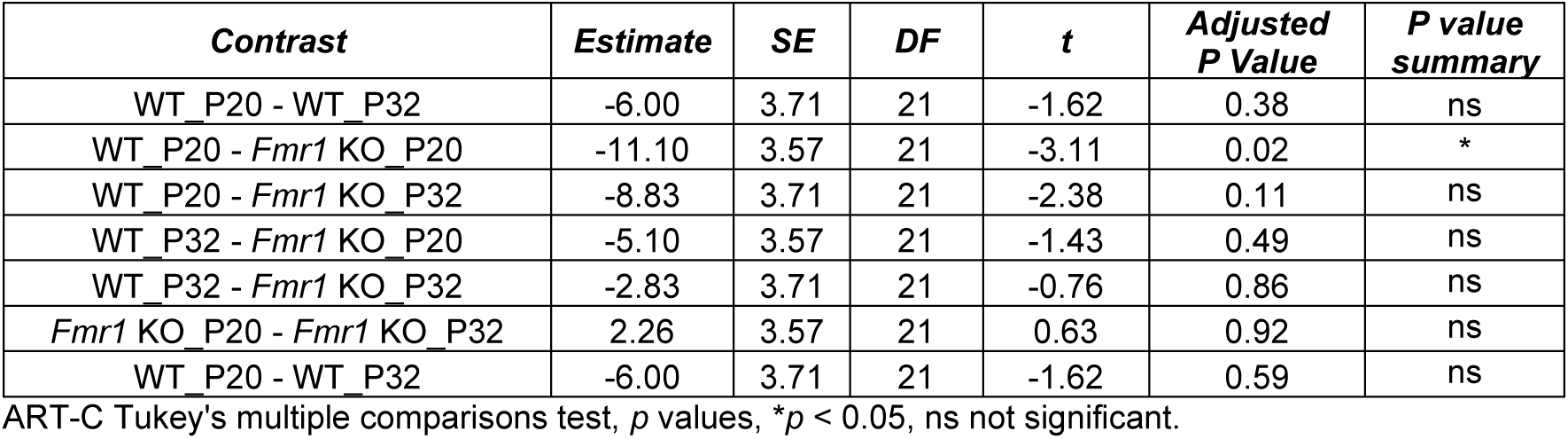
*Post hoc* pairwise of number of cFos positive cells for contrast genotype x age for mice of the following groups WT_P20 (*n*=6), *Fmr1* KO_P20 (*n*=7), WT_P32 (*n*=6), and *Fmr1 KO_P32* (*n*=6). This table is associated with data in Fig. 7B.

| Contrast | Estimate | SE | DF | t | Adjusted P Value | P value summary |
| --- | --- | --- | --- | --- | --- | --- |
| WT_P20 - WT_P32 | -6.00 | 3.71 | 21 | -1.62 | 0.38 | ns |
| WT_P20 - *Fmr1* KO_P20 | -11.10 | 3.57 | 21 | -3.11 | 0.02 | * |
| WT_P20 - *Fmr1* KO_P32 | -8.83 | 3.71 | 21 | -2.38 | 0.11 | ns |
| WT_P32 - *Fmr1* KO_P20 | -5.10 | 3.57 | 21 | -1.43 | 0.49 | ns |
| WT_P32 - *Fmr1* KO_P32 | -2.83 | 3.71 | 21 | -0.76 | 0.86 | ns |
| *Fmr1* KO_P20 - *Fmr1* KO_P32 | 2.26 | 3.57 | 21 | 0.63 | 0.92 | ns |
| WT_P20 - WT_P32 | -6.00 | 3.71 | 21 | -1.62 | 0.59 | ns |

ART-C Tukey's multiple comparisons test, $p$ values, \* $p < 0.05$ , ns not significant.

Taken together, these results suggest that *Fmr1* mutation was associated with a (prematurely) increased number of activated cells in the IC after sound exposure, especially during early auditory development.

### 3.6. AGS severity is correlated with stronger neural synchronicity in subcortical auditory generators

Epileptic seizures are caused by stereotyped changes in neurological function leading to hyper-synchronization of underlying neural networks (Medeiros and Moraes, 2014). It has been proposed that the brain pathophysiology associated with seizure susceptibility would compromise the functional synchronization of neural circuitry, even outside of the seizure state (Pinto et al., 2019; Uhlhaas and Singer, 2006), and that how easily networks synchronize to an external oscillator may predict brain state changes that lead to seizures (Pinto et al., 2017). In the auditory system, periodic oscillatory activity of neuron ensembles can be externally driven using sinusoidally amplitude-modulated sounds and measured as auditory steady-state response (ASSR) electrical potentials. ASSRs are thought to reflect the synchronous discharge of auditory neurons phase-locked to the modulation frequency of tonal stimulation (Kuwada et al., 2002; Parthasarathy and Bartlett, 2012; Picton et al., 2003). To evaluate the synchronous neural activity of subcortical auditory structures, we acoustically stimulated WT_P20, *Fmr1* KO_P20, WT_P32, and *Fmr1* KO_P32 mice with a tone (carrier frequency of 11.3 kHz) modulated in amplitude (512 Hz, Fig. 1A) and extracted the signal-to-noise ratio (SNR, maximum amplitude at the first harmonic of the modulation frequency divided by the noise floor amplitude) of the ASSR spectral magnitude (Fig. 1D). Surprisingly, we found no statistically significant main effects or interactions of genotype, sex, or age (Fig. 8A, Table 15). To assess a possible contribution of increased synchronous subcortical activity to AGS, we pooled the ASSR SNRs of *Fmr1* KO_P20 mice into four groups based on AGS phenotypes, *i.e*., *Fmr1* KO_P20 with no response, wild running, tonic-clonic seizure, or respiratory arrest. There was a significant difference in ASSR SNR between these four groups (one-way ANOVA, *p* = 0.01, *F* (3, 11) = 5.57, Fig. 8B). When compared to the *Fmr1* KO_P20 mice with no response as control (mean±SEM, 2.4±0.6), ASSR SNRs were increased significantly in mice with the most severe AGS phenotype (respiratory arrest, 7.5±1.5, Fig. 8B, Table 16), and by a statistical trend in mice with tonic-clonic seizures (6.5±1.4, Fig. 8B, Table 16), whereas those from mice with wild running were similar (2.3±0.3, Fig. 8B, Table 16). Furthermore, there was a strong positive, statistically significant, correlation between the AGS phenotypes and ASSR SNRs (Fig. 8B, Spearman r 0.7713, two-tailed *p* = 0.001). In order to assess how well the ASSR SNR predicted the AGS phenotype, we conducted a binary logistic regression test (ASSR SNR versus ‘Any seizure response (0=No/1=Yes)’). We found a statistically significant relationship between the ASSR SNR and AGS test outcome (Fig. 8C, Likelihood ratio test *p*=0.0179, Tjur’s R squared 0.3064), where the probability of any seizure response (wild running, tonic-clonic seizure, and respiratory arrest) is 50% at an ASSR SNR value of 2.39 and 75% at an ASSR SNR value of 3.61. In conclusion, these findings suggest that increased phase-locking and synchronization in the subcortical auditory processing circuits assessed through ASSRs might be associated with the predisposition to more intense AGS severity in *Fmr1* KO mice.

**Fig. 8.**
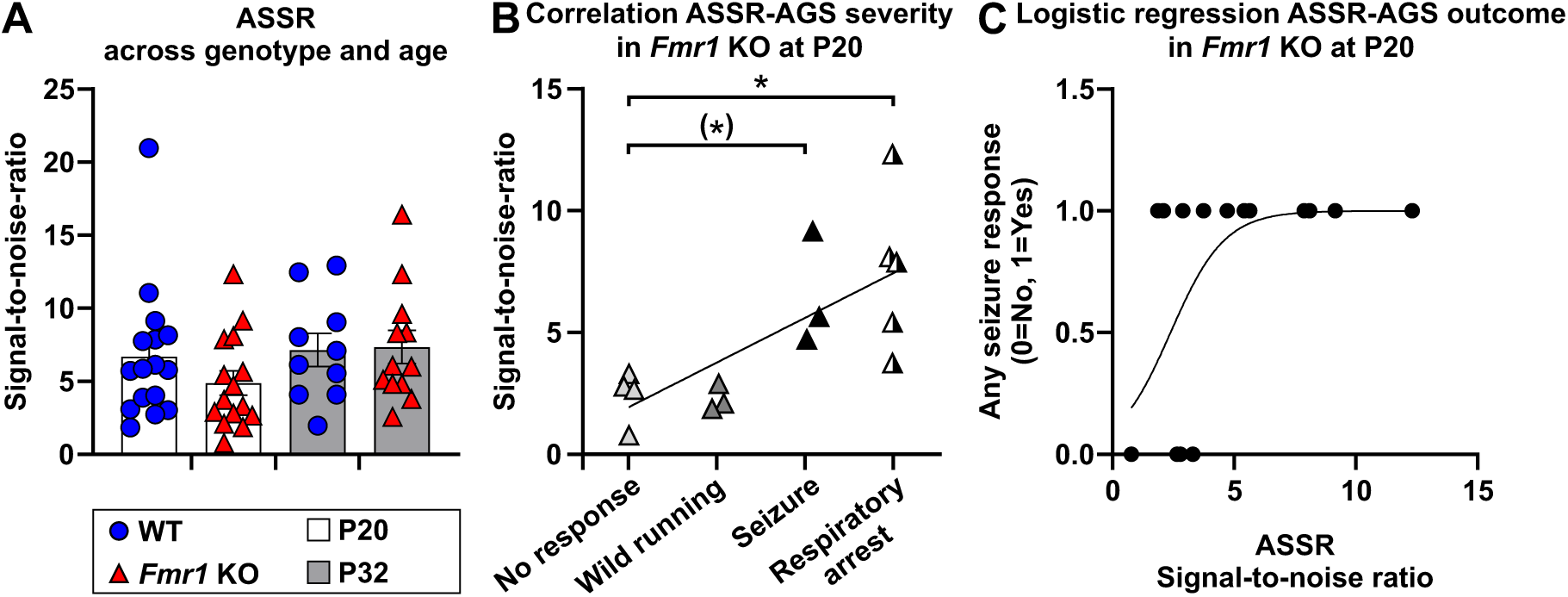
AGS severity is correlated with ASSR SNR amplitude. **(A)** ASSR SNRs from infant (white bar) and juvenile (gray bar) WT (blue circles) and *Fmr1* KO mice (red triangles) as a measure for neuronal synchronization to an external oscillator (i.e., modulated tone). ASSR SNRs were not different between WT_P20 (6.7±1.1), *Fmr1* KO_P20 (4.9±0.8), WT_P32 (7.1±1.1), and *Fmr1* KO_P32 (7.4±1.1) mice. **(B)** In infant *Fmr1* KO mice pooled by AGS phenotype, ASSR SNRs were significantly different (1-way ANOVA *p* = 0.01, *F* (3, 11) = 5.57), in particular between mice with no response and respiratory arrest, and a statistical trend in mice with tonic-clonic seizures. There was a strong positive, statistically significant, correlation between ASSR SNR and AGS severity (Spearman r 0.7713, two-tailed *p* = 0.001). **(C)** Binary logistic regression for ASSR SNR and AGS outcome grouped into no response (‘Any seizure response (0=No)’) and any AGS response including wild running, seizure, and respiratory arrest (‘Any seizure response (1=Yes)’). There was a statistically significant relationship between the ASSR SNR and AGS test outcome (Likelihood ratio test *p*=0.0179, Tjur’s R squared 0.3064, curve equation log odds = −2.145+0.8983*X). **(A)** WT_P20 (*n*=16), *Fmr1* KO_P20 (*n*=15), WT_P32 (*n*=10), *Fmr1 KO_P32* (*n*=12), **(B)** *Fmr1* KO_P20 no response (*n*=4), KO_P20 wild running (*n*=3), KO_P20 seizure (*n*=3), KO_P20 respiratory arrest (*n*=5), **(C)** ‘Any seizure response (0=No)’ (*n*=4), ‘Any seizure response (1=Yes)’ (*n*=11). Data expressed as **(A)** mean (bars) ± SEM (error bars) and individual animals (symbols), **(B)** individual animals (symbols) and linear regression line for visual clarity, or **(C)** individual animals (symbols) and binary simple logistic regression curve. *p* values, (*)p < 0.1, *p < 0.05.

**Table 15.** Statistical comparisons of ASSR SNR from mice of the following four groups: WT_P20 (*n*=16), *Fmr1* KO_P20 (*n*=15), WT_P32 (*n*=10), and *Fmr1 KO_P32* (*n*=12). This table is associated with data in Fig. 8A.

| Parameter | Source of Variation | P value | P value summary | F (DFn, DFd) | Partial $\eta^2$ |
| --- | --- | --- | --- | --- | --- |
| ASSR | Genotype | 0.37 | ns | $F(1, 45) = 0.78$ | 0.017 |
| | Sex | 0.77 | ns | $F(1, 45) = 0.08$ | 0.002 |
| | Age | 0.14 | ns | $F(1, 45) = 2.24$ | 0.047 |
| | Genotype:Sex | 0.22 | ns | $F(1, 45) = 1.51$ | 0.033 |
| | Genotype:Age | 0.37 | ns | $F(1, 45) = 0.80$ | 0.018 |
| | Sex:Age | 0.67 | ns | $F(1, 45) = 0.18$ | 0.004 |
| | Genotype:Sex:Age | 0.35 | ns | $F(1, 45) = 0.86$ | 0.019 |

3-way ART ANOVA, $p$ values, ns not significant.

**Table 16.** *Post hoc* pairwise comparisons of ASSR SNR from *Fmr1* KO_P20 mice of the following four AGS categories: no response (*n*=4), wild running (*n*=3), seizure (*n*=3), or respiratory arrest (*n*=5). This table is associated with data in Fig. 8B.

| *Parameter* | *Contrast* | *Mean Diff.* | *95% CI of diff.* | *DF* | *q* | *Adjusted P Value* | *P value summary* |
| --- | --- | --- | --- | --- | --- | --- | --- |
| ASSR | no response vs. wild running | 0.09554 | -4.660 to 4.851 | 11 | 0.05 | >0.9 | ns |
|  | no response vs. seizure | -4.126 | -8.881 to 0.6294 | 11 | 2.36 | 0.09 | ns |
|  | no response vs. respiratory arrest | -5.113 | -9.289 to -0.9359 | 11 | 3.33 | 0.01 | * |

Dunnett's multiple comparisons test, $p$ values, \* $p < 0.05$ , \*\* $p < 0.01$ , \*\*\* $p < 0.001$ , ns not significant.

### 3.7. Conclusion

This study highlights the contribution of *Fmr1* mutation to altered auditory maturation and sound responsiveness in distinct parts of the auditory pathway, during a time period that coincides with a critical window for auditory development. At the beginning of this window, i.e., in P20 mice, *Fmr1* mutation led to increased ABR wave III and IV amplitudes, generated by the SOC or LL and IC, respectively (Fig. 9A, B). The hyperresponsiveness at the level of the SOC in the brainstem appeared to be relayed to the IC in the midbrain where it was then amplified, likely facilitating these animals’ susceptibility to AGS in response to loud sounds. Towards the end of the critical window, i.e., by P32, ABR wave III and IV amplitudes had stabilized at values similar to those of WT mice (Fig. 9C), and AGS responses had ceased. Taken together, this suggests that the proper maturation of ABR waveforms, reflecting the propagating sound response along the auditory pathway, was disturbed early during this critical time window, likely contributing to FXS-like sound hypersensitivity. Furthermore, at P20, the ASSRs to amplitude-modulated stimuli, which assess periodic electrical oscillations in the subcortical auditory regions, increased in *Fmr1* KO mice as the severity of their AGS phenotype increased (Fig. 9D). This points to the notion that hyper-synchronization of subcortical auditory neuron ensembles may be linked to increased proneness to sound hypersensitivity.

**Fig. 9.**
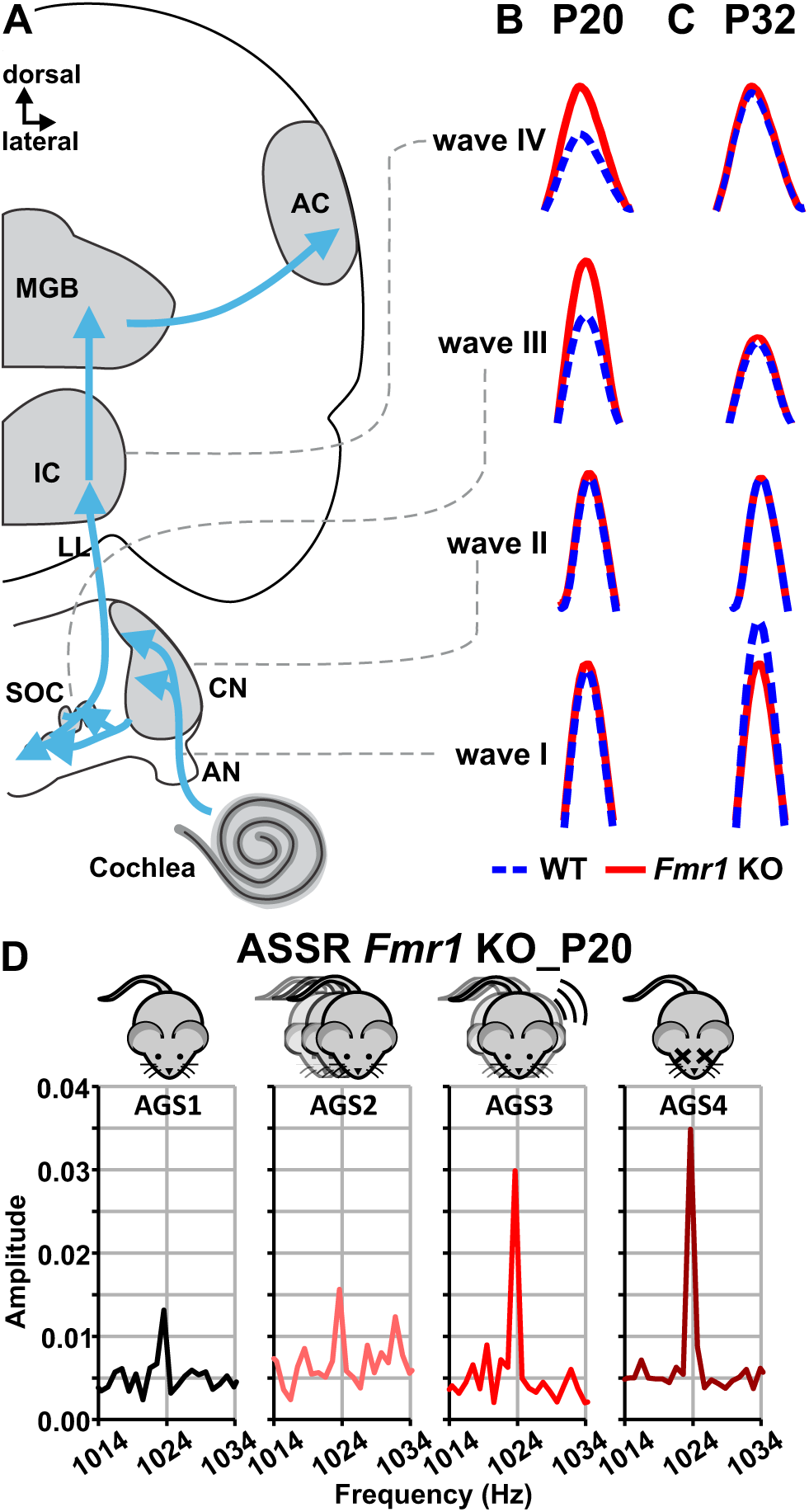
Summary of ABRs in infant (P20) and juvenile (P32) WT and *Fmr1* KO mice, and of ASSRs in the four AGS phenotypes of *Fmr1* KO_P20 mice. (A) Schematic drawing of the auditory pathway and correlated stimulus-evoked deflections of ABR waves, I, II, III, and IV. The auditory signal along the auditory pathway measured by ABRs provides information regarding auditory function and hearing sensitivity. The different peaks of the ABR wave can be assigned to distinct parts of the ascending auditory pathway: Wave I is generated by the AN, wave II by the CN, wave III by the SOC, and wave IV by the LL and IC. (B) *Fmr1* KO_P20 mice (red solid line) had increased ABR wave III and IV amplitudes compared with WT_P20 mice (blue dashed line). In contrast to infants, **(C)** ABR wave III and IV amplitudes were similar in *Fmr1* KO_P32 mice (red solid line) and WT_P32 mice (blue line). ABR wave I amplitudes remained lower in *Fmr1* KO_P32 than in WT_P32 mice. **(D)** ASSRs amplitudes as a measure for neuronal synchronization phase-locked to the modulation frequency, pooled and averaged for *Fmr1* KO_P20 mice depending on their AGS severity: no response (AGS1, black line), wild running (AGS2, light red line), tonic-clonic seizure (AGS3, red line), respiratory arrest (AGS4, dark red line). ASSR magnitudes increased in *Fmr1* KO_P20 mice with AGS severity. Abbreviations AC, auditory cortex; AGS, AGS; AGS1, no response; AGS2, wild running; AGS3, tonic-clonic seizure; AGS4, respiratory arrest; AN, auditory nerve; CN, cochlear nucleus; IC, inferior colliculus; LL, lateral lemniscus; MGB, medial geniculate body; SOC, superior olivary complex.

In summary, our findings indicate that higher neural responsiveness to sound, as well as hyper-synchronization to a continuous modulated sound, promote susceptibility to and severity of AGS in infant *Fmr1* KO mice.

## 4. Discussion

In this study, we report that *Fmr1* mutation leads to characteristic changes in sound processing from auditory periphery to midbrain, likely due to altered maturation of the auditory system during early postnatal development. Notably, our study includes both male and female mice, which is an important distinction as previous studies have often focused predominantly on males. The atypical central, but not so much peripheral, sound processing appeared to contribute to the susceptibility and severity of AGS, modelling auditory hypersensitivity in FXS.

### 4.1. Altered maturation of the auditory periphery does not directly underlie AGS susceptibility

The auditory periphery represents the initial stages of auditory processing and includes the cochlear sensory epithelium and the auditory nerve. These structures can be assessed for their functional integrity through ABR thresholds and ABR wave I, respectively. Auditory thresholds are known to provide a sensitive measure of outer hair cell (OHC) function. These cells in the cochlear sensory epithelium play a crucial role in cochlear amplification, which enhances hearing sensitivity (Dallos, 2008; Dallos and Evans, 1995; Liberman and Beil, 1979; Liberman and Kiang, 1978). The maturation of the cochlea follows what is known as the “sliding place principle” (Rubel et al., 1984), i.e., from middle-to-apical and middle-to-basal cochlear regions. Accordingly, the developmental increase in hearing sensitivity is frequency-dependent (Ehret, 1976). In the present study, we found no significant differences between the ABR thresholds in WT and *Fmr1* KO mice at P20 for 4 kHz and 11 kHz (Fig. 3), suggesting WT-like maturation in the *Fmr1* KO mice by that age at these low to mid frequencies, represented in the apical and medial cochlear regions. However, at 32 kHz, *Fmr1* KO mice showed higher ABR thresholds (Fig. 3), indicative of delayed cochlear maturation of the basal sensory epithelium. A recent study suggested that FMRP is involved in the development of the auditory periphery, with distinct developmental trajectories across cell types. In the cochlea, FMRP is expressed in the hair cells of the sensory epithelium during the prehearing period (hearing onset in mice is ∼P12, Mikaelian and Ruben, 1965), and is then strongly downregulated afterward during maturation and in young adults (3 months old), with no discernible tonotopic variation in cellular FMRP intensity (Wang et al., 2023). Consistent with the idea that FMRP plays a critical role in hearing onset (Yu and Wang, 2023) and timely cochlear maturation rather than in maintaining mature cochlear function, juvenile *Fmr1* KO mice in the present study (Fig. 3), as well as young adult *Fmr1* KO mice (Ferraguto et al., 2023), showed similar ABR thresholds to WT controls across low to high frequencies. This suggests that while cochlear maturation is delayed in the absence of FMRP, ABR thresholds eventually stabilize at normal levels at a time when FMRP expression would naturally be low in WT mice. In addition to ABR thresholds, Ferraguto et al. (2023) assessed OHC function through distortion product otoacoustic emissions, which were unaltered in 3-month-old *Fmr1* KO mice. Taken together, these findings suggest that in absence of FMRP expression, cochlear maturation occurs in the proper tonotopic order but follows a delayed trajectory, and that auditory thresholds reach WT-like levels by the juvenile stage and remain stable at least until young adulthood.

In contrast to FMRP expression in cochlear hair cells, FMRP in spiral ganglion neurons, i.e. cell bodies of auditory nerve fibers, is strong during prehearing and further upregulated with maturation and across age (Wang et al., 2023). Studies have consistently shown that *Fmr1* KO mice exhibit little to no change in ABR wave I amplitudes, as a measure for the summed sound-evoked activity of the auditory nerve, during early postnatal development (P15, El-Hassar et al., 2019; P20, Fig. 5C of the present study) and diminished wave I amplitudes in the juvenile stage (P32, Fig. 5D of the present study) and adulthood (Ferraguto et al., 2023; Rotschafer et al., 2015). Ferraguto et al. (2023) showed that the lower wave I amplitude in *Fmr1* KO mice is accompanied by a loss of ribbon synapses of inner hair cells with dendrites from spiral ganglion neurons of medial cochlear turns, indicating cochlear deafferentation (Kujawa and Liberman, 2009). Such a lack of active-zone-anchored synaptic ribbons reduces the presynaptic readily releasable vesicle pool, and impairs the synchronous sound-evoked activation of spiral ganglion neurons (Khimich et al., 2005). Glutamatergic transmission in the spiral ganglion neurons is crucial for activity-dependent maturation and neuronal diversification of hearing function (Petitpré et al., 2022; Petitpré et al., 2018; Shrestha et al., 2018; Sun et al., 2018), in particular fast excitatory synaptic transmission of cochlear sound encoding via unique α-amino-3-hydroxy-5-methyl-4-isoxazole propionic acid receptors (AMPARs) (Gardner et al., 1999; Glowatzki and Fuchs, 2002; Raman et al., 1994; Ruel et al., 1999). Beneath inner hair cells, the AMPAR subunits GluA2/3 remain stable throughout development and into adulthood, while synaptic GluA4-containing AMPARs are upregulated in adulthood (but not populated with GluA1, Huang et al., 2012; Matsubara et al., 1996; Niedzielski and Wenthold, 1995; Parks, 2000). This upregulation may strengthen ribbon synapses with auditory nerve fibers in response to sound stimulation (Huang et al., 2012). While more studies seem to be needed to address how absence of FMRP affects GluA2, GluA3, and GluA4 in the auditory periphery, it is known that in hippocampal and frontal brain tissue lack of FMRP leads to impaired activity-dependent synaptic shuttling of AMPAR subunits between the cell surface and intracellular compartments (Chojnacka et al., 2023). Dysregulation of GluA2 and GluA4 subunit relative abundance and alterations in pre- and postsynaptic ultrastructure at the cochlear ribbon synapses in GluA3 KO mice have pathological, presymptomatic effects (Rutherford et al., 2023) and precede reduction in ABR wave I amplitudes in young adults (García-Hernández et al., 2017). Taken together, it appears possible that FMRP is involved in the regulation of AMPAR subunit composition and trafficking at the cochlear ribbon synapses, and its absence may disrupt the activity-dependent maturation of auditory synaptic transmission, contributing to early pathological changes that precede reductions in auditory nerve sound responses in juvenile *Fmr1* KO mice in the present study (Fig. 5D).

Interestingly, neither altered maturation of ABR thresholds nor wave I amplitudes was directly correlated with AGS susceptibility. Therefore, our findings support the notion that auditory hypersensitivity in FXS is due to a central, rather than peripheral, dysfunction of the auditory pathway in both the mouse model (El-Hassar et al., 2019; Garcia-Pino et al., 2017) and humans (Arinami et al., 1988; Castrén et al., 2003; Ferri, 1989; St Clair et al., 1987; Van der Molen et al., 2012).

It is important to note that altered development of the auditory periphery is likely to have secondary effects on central auditory maturation and processing, since a peripheral influence on the development and function of the central auditory system is well-documented (Lesicko and Llano, 2017; Polley et al., 2013; Rubel and Fritzsch, 2002; Ryugo, 2015). Conditional *Fmr1* KO mice with selective deletion of FMRP in spiral ganglion neurons, but not in ventral CN neurons, recapitulate the delayed closure of a critical period for neuronal loss in the ventral CN observed in constitutive *Fmr1* KO mice (Yu and Wang, 2023). This delay temporally coincides with reduced hearing sensitivity, suggesting an association between auditory brain development and sensory inputs shaped by peripheral FMRP (Yu and Wang, 2023). Future studies should investigate to what extent correcting the maturation of the auditory periphery can rectify central auditory function and hypersensitivity in *Fmr1* KO mice.

### 4.2. Correlation between altered maturation of the auditory brainstem and AGS susceptibility

In patients with FXS, auditory processing impairment is predominantly evident in the prolonged latencies of ABR waves III and V (equivalent to rodent wave IV) and in the interpeak intervals III-V and I-V (Arinami et al., 1988). Consistent with these findings, the increased amplitudes of ABR waves III (present study, Fig.5C) and IV (P15, El-Hassar et al., 2019; P20, Fig. 5C of the present study) in infant *Fmr1* KO mice suggest altered excitability of brainstem nuclei as the underlying neuronal basis for auditory hypersensitivity, rather than peripheral dysfunction. Amplitudes for peaks II, III, and IV in the juvenile stage (Fig. 5D of the present study) and adulthood (12 kHz, Rotschafer et al., 2015) were unaffected, when AGS susceptibility had subsided. Few or no genotype differences were found for peak latencies of ABR waves I, II, III, and IV across early (Fig. 4 of the present study; I and IV, El-Hassar et al., 2019) or late development (Fig. 4 of the present study) and adulthood (Chawla and McCullagh, 2022; Rotschafer et al., 2015). As discussed in Rotschafer et al. (2015), the absence of a latency effect could be due to the smaller size of the mouse auditory pathway, differences in auditory processing between species, or varying responses to mutations in *Fmr1*.

Analyzing the slopes of ABR amplitude growth functions, we found an increased sensitivity of the neural response to increasing sound intensity with AGS susceptibility, particularly on the level of the LL and IC (Fig. 6A). Single unit recordings have previously shown that IC neurons from *Fmr1* KO mice during development produce increased response magnitudes to tone bursts and amplitude-modulated tones and have broader frequency tuning curves compared with age-matched WT neurons (Nguyen et al., 2020), the latter suggesting that more IC neurons respond synchronously to sounds (Rotschafer and Razak, 2013). Using immunostaining for cFos to determine the number of activated cells, Nguyen et al. (2020) demonstrated that a genotype difference between *Fmr1* KO and WT mice in cFos-positive cell density was evident in the dorsolateral half of the IC at P21 and shifted to the ventromedial half of the IC at P34. In the same study, to avoid inducing AGS and minimize the potential confounding effects of locomotor activity (Yang et al., 2020), sounds at 85 dB SPL were used for infants, and 80- or 90-dB SPL for juveniles. In our experiments, the same acoustic stimulation across all groups (∼90 dB SPL) did not show any age-related effects on cFos-positive cell density throughout the entire dorso-ventral axis of the IC (Fig. 7). The genotype effect was primarily driven by a higher number of cFos-labeled cells in *Fmr1* KO mice at P20 compared to age-matched WT controls (Fig. 7). We also did not observe a systematic influence of the AGS phenotype on cFos-positive cell counts, though a higher sample size is needed to substantiate these claims. It is worth noting that the sound exposure in Nguyen et al. (2020) and our study differed in frequency range and sound modulation, possibly contributing to the slightly different outcomes in cFos cell counts. Taken together, these findings suggest that the level of activity in the IC may correlate with auditory hypersensitivity, particularly during early auditory development. Interestingly, there was no evidence for abnormal development of IC frequency tonotopy (Nguyen et al., 2020), matching our respective conclusion on the auditory periphery.

Gonzalez et al. (2019) have shown that *Fmr1* deletion in glutamatergic neurons of the IC is necessary, but not sufficient, to induce AGS. While *Fmr1* deletion in these neurons alone does not fully recapitulate the AGS phenotype, it is required in conjunction with *Fmr1* deletion in other subcortical vGlut2 (vesicular glutamate transporter 2)-expressing glutamatergic neurons, to cause AGSs (Gonzalez et al., 2019). Previous data suggests that the most likely subcortical sources of vGlut2 terminals in the IC are the IC itself, the intermediate nucleus of the LL, the dorsal CN, and the medial and lateral superior olive (MSO, LSO) (Ito and Oliver, 2010). In the LSO, vGlut2 is the dominant vGlut (Blaesse et al., 2005) and appears to be obligatory in glutamatergic LSO neurons (Ito et al., 2011). Between prehearing (P6) and shortly after hearing onset (P14), fractional coverage of vGlut2 expression in the LSO decreases in both WT and *Fmr1* KO mice (Rotschafer and Cramer, 2017), however, *Fmr1* KO mice go on to show greater vGlut2-immunolabelled area within the LSO (Rotschafer et al., 2015). In line with this morphological difference, during the first 10 days after hearing onset, the functional maturation of synapses in the LSO is severely altered in *Fmr1* KO mice, resulting in enhanced excitatory input strength from the ipsilateral side and an excessive number of excitatory synapses (Garcia-Pino et al., 2017). LSO neurons in these mice exhibit enhanced firing rates, broadened tuning curves and shifted interaural level differences (Garcia-Pino et al., 2017), which are one of the major cues for localizing sound in the horizontal plane (Grothe et al., 2010). Additionally, morphological alterations of the SOC nuclei and neurons have been observed in post-mortem tissue of subjects with FXS and autism (Kulesza et al., 2011; Kulesza and Mangunay, 2008). In the present study, while the slopes of ABR wave III amplitude growth functions were not significantly different between AGS susceptibility phenotypes (Fig. 6A), we found a strong correlation between the slopes of waves III and IV (Fig. 6B). These findings emphasize that auditory hyperresponsiveness underlying AGS may, at least in part, originate in the SOC and then be relayed to and pathologically amplified in the IC of the auditory midbrain. Investigating the pathophysiological mechanisms underlying these auditory processing abnormalities in the SOC and IC may be key to understanding brainstem circuit-level alterations that lead to auditory hypersensitivity in FXS and autism.

### 4.3. ASSR amplitudes might be useful to predict AGS severity

We found no differences in ASSR SNR phase-locked to amplitude-modulated sounds between WT and *Fmr1* KO mice at P20 and P32 from subcortical auditory neurons (Fig. 8A). This is consistent with Nguyen et al. (2020), who found no differences in single neuron phase locking to amplitude-modulated tones in electrophysiological recordings from the IC between these genotypes at P14, P21, and P34. During maturation, the envelope following response to amplitude modulated tones increases with age in rats (Prado-Gutierrez et al., 2012; Venkataraman and Bartlett, 2014) and humans (Pethe et al., 2004; Savio et al., 2001; Mijares Nodarse et al., 2012), and the modulation frequency at which the maximum amplitude can be obtained is shifted to higher values depending on the carrier frequency (Prado-Gutierrez et al., 2012). It is possible that age-related and/or genotypic differences would have become apparent with a more comprehensive combination of carrier and modulation frequencies.

When pooled for AGS severity (no response, wild running, seizure, or respiratory arrest), we found a strong positive correlation between the AGS phenotypes and ASSR SNRs (Fig. 8B), as well as a significant relationship between the ASSR SNR and AGS test outcome in *Fmr1* KO mice at P20 (Fig 8C). In line with our findings, Wistar audiogenic rats show increased ASSR-normalized energy even when operating within sound stimulation intensities that are subthreshold to AGS (85 dB SPL, Pinto et al., 2017). These findings suggest that ASSR could be a valuable tool for assessing the susceptibility of brain networks to hyper-synchronization, potentially serving as a diagnostic tool and even predicting brain state shifts that lead to AGS. Interestingly, in humans, the ASSR amplitude function is a good neural correlate of the loudness growth function (e.g., inaudible to unbearable, Van Eeckhoutte et al., 2016), speaking to the usefulness of ASSR amplitudes as an electrophysiological correlate to estimate the loudness percept and possibly auditory hypersensitivity. Future work is necessary to determine the mechanisms that underlie the altered phase-locking, in particular depending on AGS severity. In the aforementioned study by Prado-Gutierrez et al. (2012), the authors concluded that the increase in the envelope following response during the two weeks after hearing onset is due to the increased sensitivity of the auditory system during maturation (Prado-Gutierrez et al., 2012), and could be a consequence of axonal myelination, the maturation of the synaptic mechanisms involved in the response, and the progressive decrease of the input membrane resistance and the membrane time-constant of the neuron. These developmental processes would contribute to an increased capacity of neurons to follow high-frequency stimulation and enhance the synchrony of neural responses to stimuli, thereby producing larger response amplitude (Ryugo et al., 2006; Scott et al., 2005). Another potential hypothesis for the observed differences in ASSR and AGS activity in *Fmr1* KO mice could be that some of these animals may be more vulnerable to stress. In absence of FMRP, stress produces more immediate increases in corticosteroids (Ghilan et al., 2015), which might in turn affect ASSR magnitudes (Marchetta et al., 2022).

The ASSR can be dominated by cortical (Herdman et al., 2002a; Herdman et al., 2002b; Kuwada et al., 2002) and subcortical (Parthasarathy and Bartlett, 2012; Picton et al., 2003) sources, depending on the specific modulation frequency used to elicit the response. Early neural sources dominate amplitude modulation ranges at ∼1000 Hz, which is close to the peak phase-locking capacity of the auditory nerve (Joris and Yin, 1992; Palmer and Evans, 1982; Shaheen et al., 2015) and temporal properties of the primary cochlear nucleus neurons (Joris et al., 2004). Generators of responses to the 200–500 Hz amplitude modulation range are thought to be primarily in the brainstem and midbrain (Kiren et al., 1994; Kuwada et al., 2002; Parthasarathy and Bartlett, 2012). In conclusion, while no differences in ASSR SNR phase-locking were found between WT and *Fmr1* KO mice, the relationship between ASSR magnitudes and AGS severity suggests that ASSR could serve as a useful tool for assessing altered neural processing in specific neuronal generators that contributes auditory hypersensitivity.

### 4.4. Limitations

While *Fmr1* KO mice have significantly advanced our understanding of FXS, it is crucial to acknowledge that these models do not fully replicate the human condition. In humans, a full *FMR1* gene mutation contains over 200 CGG repeats, leading to promoter hypermethylation and gene silencing. This reduces levels of the *FMR1* gene product, FMRP, central to FXS pathology (Hagerman et al., 2001; Oberlé et al., 1991; Verkerk et al., 1991). In contrast to humans, the *Fmr1* KO mouse, which is the most well characterized rodent model, lacks FMRP protein due to a disruption in its *Fmr1* gene (Bakker et al., 1994; Yan et al., 2004). Such mechanistic differences might underlie the findings that the *Fmr1* KO mouse replicates certain features of FXS (such as neuronal morphology and hyperactivity), while other clinical features are not reproduced or inconsistent (such as cognitive impairments and anxiety) (reviewed in e.g., Kazdoba et al., 2014; Willemsen and Kooy, 2023). Phenotypes can also depend on the background strain (e.g., Pietropaolo et al., 2011). In terms of sensory hypersensitivity, *Fmr1* KO rodent models generally align with observations in humans with FXS. These models exhibit behaviors such as seizures, abnormal visual-evoked responses, auditory hypersensitivity, and disrupted processing at various levels of the auditory system. This suggests that fundamental sensory processing circuits are likely conserved across species, providing a valuable translational platform for identifying biomarkers, understanding underlying mechanisms, and evaluating potential drug targets to alleviate sensory symptoms associated with FXS (reviewed in e.g., Rais et al., 2018). Furthermore, in order to assess acute temporal dynamics of ASSR underlying AGS susceptibility and severity, specifically in the pre-ictal, ictal, and post-ictal period, measurements in awake animals might be beneficial.

### 4.5. Conclusion

This study highlights that *Fmr1* mutation causes disruptions in auditory processing, primarily in the central auditory system, rather than the auditory periphery, and these changes contribute to auditory hypersensitivity in FXS. Specifically, delayed maturation of cochlear and auditory nerve function does not directly correlate with AGS susceptibility. Instead, altered central processing, especially in the brainstem and midbrain, appears to be the primary driver of AGS. Finally, our findings suggests that early developmental interventions targeting auditory processing might offer a critical therapeutic window to alleviate auditory hypersensitivity in FXS.

## Abbreviations

a: cerebral aqueduct
ABR: Auditory brainstem response
AC: Auditory cortex
AGS: audiogenic seizure
AGS1: No response
AGS2: Wild running
AGS3: Tonic-clonic seizure
AGS4: Respiratory arrest
AN: Auditory nerve
ART: Aligned rank transform
ART-C: ART contrast tests
ASSR: Auditory steady state response
CB: Cerebellum
CF: Carrier frequency
CUN: Cuneiform nucleus
CN: Cochlear nucleus
DCN: Dorsal cochlear nucleus
FFT: Fast Fourier transformation
*Fmr1*: Fragile X messenger ribonucleoprotein 1
FMRP: Fragile X messenger ribonucleoprotein 1
FXS: Fragile X syndrome
HB: Hindbrain
IC: Inferior colliculus
ICc: Inferior colliculus central nucleus
ICd: Inferior colliculus dorsal nucleus
ICe: Inferior colliculus, external nucleus
IQR: Interquartile range
KO: Knockout
LSO: Lateral superior olive
LL: Lateral lemniscus
m: Medial
MF: Modulation frequency
MGB: Medial geniculate body
MSO: Medial superior olive
OHC: Outer hair cell
PAG: Periaqueductal gray
P: Postnatal day
SEM: Standard error of the mean
SNR: Signal-to-noise ratio
SOC: Superior olivary complex
vGlut2: Vesicular glutamate transporter 2
VCN: Ventral cochlear nucleus
WT: Wildtype
v: Ventral.

## Author contributions

DMö designed the study, performed experiments, MATLAB coding, formal analysis, data interpretation and visualization, and wrote the original draft. DMa performed experiments, formal analysis, and participated in writing the original draft. WX participated in experiments and editing of the manuscript. JY provided resources and contributed to editing of the manuscript. NC provided funding, contributed to study design and editing of the manuscript. All authors read and approved the final manuscript.

## Data availability

Data will be made available on request.

