## Supplementary Figure 1 for "Altered auditory maturation in Fragile X syndrome and its involvement in audiogenic seizure susceptibility"

### *Supplementary Material*

#### 1 Supplementary Figures

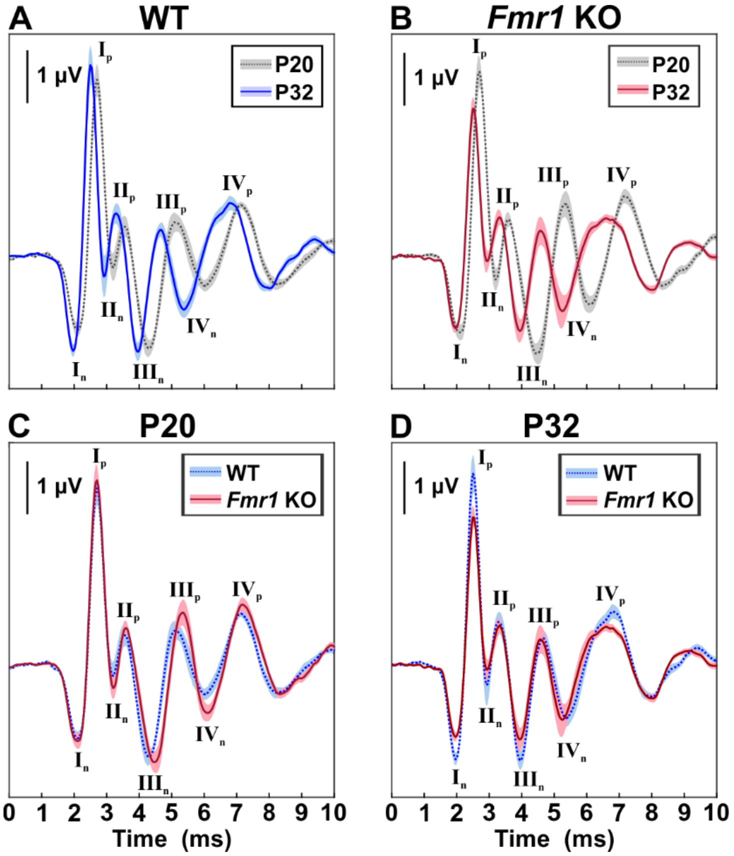


**Supplementary Figure 1. ABR waveforms at 60 dB *re* threshold.** ABR waveforms consist of consecutive amplitude deflections (waves), with each wave consisting of a starting negative (n) peak and the following positive (p) peak. ABR wave I: $I_n$-$I_p$, wave II: $II_n$-$II_p$, wave III: $III_n$-$III_p$, wave IV: $IV_n$-$IV_p$. Wave latencies were defined by the onset timing (negative peak) of each corresponding wave, and wave amplitudes by the peak-to-peak difference. ABR waveforms for **(A)** WT_P20 (gray dotted line and area) and WT_P32 (blue line and area), **(B)** *Fmr1* KO_P20 (gray dotted line and area) and *Fmr1* KO _P32 (red line and area), **(C)** WT_P20 (blue dotted line and area) and *Fmr1* KO_P20 (red line and area), and **(D)** WT_P32 (blue dotted line and area) and *Fmr1* KO_P32 (red line and area). WT_P20 (*n*=16), *Fmr1* KO_P20 (*n*=15), WT_P32 (*n*=10), and *Fmr1* KO_P32 (*n*=12). Data expressed as mean (lines) ± SEM (shaded areas).
