## Supplementary Table 1 for "Altered auditory maturation in Fragile X syndrome and its involvement in audiogenic seizure susceptibility"

### 2 Supplementary Tables

**Supplementary Table 1.** Statistical comparisons of ABR thresholds from mice of the following four groups: WT_P20 (*n*=16), *Fmr1* KO_P20 (*n*=14-15), WT_P32 (*n*=10), and *Fmr1 KO_P32* (*n*=12). The thresholds for one *Fmr1* KO_P20 animal at 4 and 32 kHz were excluded from analysis because the acoustic stimuli were presented in 10 dB instead of 5 dB steps.

| ***Parameter*** | ***Source of Variation*** | ***P value*** | ***P value summary*** | ***F (DFn, DFd)*** | ***ηp2*** |
|---|---|---|---|---|---|
| ABR threshold | Genotype | 0.31 | ns | F(1, 44.59) = 1.02 | 0.022 |
| | Sex | 0.29 | ns | F(1, 44.59) = 1.10 | 0.024 |
| | Age | <0.001 | *** | F(1, 44.54) = 22.76 | 0.338 |
| | Frequency | <0.001 | *** | F(2, 88.49) = 398.98 | 0.900 |
| | Genotype:Sex | 0.60 | ns | F(1, 44.58) = 0.26 | 0.006 |
| | Genotype:Age | 0.36 | ns | F(1, 44.58) = 0.83 | 0.018 |
| | Sex:Age | 0.06 | ns | F(1, 44.56) = 3.54 | 0.074 |
| | Genotype:Frequency | 0.01 | * | F(2, 88.53) = 4.70 | 0.096 |
| | Sex:Frequency | 0.20 | ns | F(2, 88.53) = 1.59 | 0.035 |
| | Age:Frequency | 0.01 | * | F(2, 88.53) = 4.11 | 0.085 |
| | Genotype:Sex:Age | 0.21 | ns | F(1, 44.59) = 1.58 | 0.034 |
| | Genotype:Sex:Frequency | 0.44 | ns | F(2, 88.52) = 0.82 | 0.018 |
| | Genotype:Age:Frequency | 0.62 | ns | F(2, 88.53) = 0.46 | 0.010 |
| | Sex:Age:Frequency | 0.73 | ns | F(2, 88.53) = 0.31 | 0.007 |
| | Genotype:Sex:Age:Frequency | 0.23 | ns | F(2, 88.54) = 1.49 | 0.033 |

Mixed effects ART ANOVA, *p* values, $*p < 0.05$, $***p < 0.001$, ns not significant.
