## Supplementary Table 2 for "Altered auditory maturation in Fragile X syndrome and its involvement in audiogenic seizure susceptibility"

### 2 Supplementary Tables

**Supplementary Table 2.** *Post hoc* pairwise comparisons of ABR thresholds for factor Frequency from mice of the following four groups: WT_P20 (*n*=16), *Fmr1* KO_P20 (*n*=14), WT_P32 (*n*=10), and *Fmr1 KO_P32* (*n*=12).

| ***Parameter*** | ***Contrast*** | ***Estimate*** | ***SE*** | ***DF*** | ***t*** | ***Adjusted P Value*** | ***P value summar*** |
|---|---|---|---|---|---|---|---|
| Frequency (kHz) | 11.3 - 32 | -45.24 | 3.54 | 88.647 | -12.78 | <0.001 | *** |
| | 11.3 - 4 | -99.89 | 3.54 | 88.647 | -28.21 | <0.001 | *** |
| | 32 - 4 | -54.65 | 3.54 | 88.190 | -15.40 | <0.001 | *** |

ART-C Tukey's multiple comparisons test, *p* values, ****p* < 0.001, ns not significant.
